# Motion Displaces Population Receptive Fields in the Direction Opposite to Motion

**DOI:** 10.1101/759183

**Authors:** Marian Schneider, Ingo Marquardt, Shubharthi Sengupta, Federico De Martino, Rainer Goebel

**Affiliations:** Faculty of Psychology and Neuroscience Maastricht University

**Keywords:** visual neuroscience, position perception, population receptive fields, visual field projections

## Abstract

Motion signals can bias the perceived position of visual stimuli. While the apparent position of a stimulus is biased in the direction of motion, electro-physiological studies have shown that the receptive field (RF) of neurons is shifted in the direction opposite to motion, at least in cats and macaque monkeys. In humans, it remains unclear how motion signals affect population RF (pRF) estimates. We addressed this question using psychophysical measurements and functional magnetic resonance imaging (fMRI) at 7 Tesla. We systematically varied two factors: the motion direction of the carrier pattern (inward, outward and flicker motion) and the contrast of the mapping stimulus (low and high stimulus contrast). We observed that while physical positions were identical across all conditions, presence of low-contrast motion, but not high-contrast motion, shifted perceived stimulus position in the direction of motion. Correspondingly, we found that pRF estimates in early visual cortex were shifted against the direction of motion for low-contrast stimuli but not for high stimulus contrast. We offer an explanation in form of a model for why apertures are perceptually shifted in the direction of motion even though pRFs shift in the opposite direction.

## 1 Introduction

An important task of the visual system is to infer the location of objects in our environment. A wide range of psychophysical studies shows that motion signals lead to systematic localisation biases [1, 2, 3, 4, 5, 6, 7, 8, 9]. In illusions called motion-induced position shifts (MIPS), a coherent motion signal shifts the apparent location of a stimulus [1]. For example, when drifting Gabor patches are presented within a stationary aperture, the stimulus appears shifted in the direction of motion [2, 6, 7]. Such illusions raise the question how our visual system encodes location and how, in the case of MIPS, the apparent position shift can be explained. Furthermore, they offer a dissociation between the physical and the perceived position of a stimulus that can clarify which neuronal processes correspond to the apparent position of the stimulus.

The magnitude of MIPS is known to depend on spatial and temporal properties of the stimulus. MIPS are larger when the stimulus is shown for longer duration (tested up to 453 ms; [6]), presented at higher speed [6, 9] or at higher eccentricities [10, 6, 9]. The magnitude of MIPS furthermore depends on spatial blurring of the presented stimulus. Blurred stimulus edges lead to larger perceptual displacements than sharp edges [4, 9] and increasing the size of the Gaussian envelope of a Gabor stimulus yields larger MIPS [4]. Arnold et al. [7] have suggested that MIPS are driven by modulation of apparent contrast of the stimulus. Supporting this suggestion, they reported perceived position shifts when observers were asked to match the extremities of two contrast envelopes (low-contrast region), but no corresponding shifts when observers were instructed to match the phase within two gratings (high-contrast region).

While the stimulus-related factors that influence MIPS are well studied, the neuronal mechanisms underlying MIPS in humans have remained controversial. Based on the results of a functional magnetic resonance imaging (fMRI) study, Whitney and colleagues [11] initially suggested that the perceptual shift was accompanied by a shift in retinotopic representation of the stimulus. They presented Gabor stimuli that contained motion drifting either towards or away from fixation. Although Gabor stimuli were presented at the same physical location, when contrasting inward and outward motion conditions, the authors observed positive signal differences in an anterior V1 region (representing higher eccentricities) and negative signal differences in a more posterior V1 region (representing lower eccentricities). Based on this observation, as well as several control experiments, they concluded that retinotopic representation is flexible and shifted by the motion direction signals of a stimulus.

This account has since been challenged by Liu and colleagues [12]. In a series of experiments, they convincingly demonstrated that retinotopic representations in V1-V3 of stimuli presented at the same location but with opposing motion are, to a large degree, overlapping. Furthermore, in one of their control experiments, they determined the preferred eccentricity for V1 voxels presenting a traveling wave stimulus that contained either inward or outward motion. Eccentricity estimates did not vary significantly between the two motion directions. Based on these results, they argued that the activation differences reported by [11] were caused by small signal differences at retinotopic locations that represent the trailing edge of motion and were amplified by contrasting motion conditions directly (since the hemodynamic response in fMRI acts like a compressive nonlinearity). Thus, although the center of mass of the retinotopic representations was slightly shifted against the direction of motion, the two representations showed substantial overlap.

In light of recent studies, the results presented by [11] and [12] can both be interpreted in terms of a phenomenon that has been termed “aperture-inward” bias [13]. [13] demonstrated that the trailing edge of a motion stimulus evokes larger responses than the leading edge, at least in early visual cortex (V1-V3) but not in V5/hMT+. Furthermore, several studies showed that responses decrease gradually with distance from the trailing edge [13, 14, 15].

Although the effects initially reported by [11] are now better understood, it still remains elusive which neuronal mechanisms underlie perceptual position shifts in humans. When monkeys and cats were presented with motion stimuli that lead to perceptual shifts for human observers, electro-physiological recordings revealed that the receptive field (RF) of V1 [10] and V4 [16] neurons were shifted against the direction of motion. Importantly, these RF shifts cannot be understood in terms of the aperture-inward bias. If response amplitudes are larger at the trailing edge of the stimulus and an RF is assumed to be composed of inputs from pre-synaptic neurons with smaller RF fields, then RFs should shift in the direction of motion, which is the opposite of what was observed empirically [10].

Using fMRI in humans it is not possible to measure RF properties of individual neurons. However, a quantitative framework for estimating the population RF (pRF) of voxels has been developed [17]. Using this framework, Harvey et al. [18] reported that pRF size and eccentricity depended on the speed and direction of a motion stimulus. In particular, higher speed led to higher pRF size and eccentricity estimates in early- and mid-level visual areas if motion was in the direction of the pRF measurement but not if motion was in an orthogonal direction. However, since opposing motion directions were presented during the same stimulus block, it was not possible to disentangle the effects of opposing motion directions on pRF eccentricity and size. Consequently, it is still unclear whether pRFs are displaced with motion in humans (and, if so, in which direction).

As a result, there is an unsatisfactory gap between human psychophysics, animal electro-physiology, and human neuroimaging. In an attempt to bridge this gap, we identified several factors that could explain the mismatch between results in human and non-human subjects. These factors motivated the the current study. First, to determine preferred eccentricity for V1 voxels, [12] presented ring apertures, while we decided to employ annulus sector apertures instead. This difference might be important since the two aperture types lead to perceptually different phenomena: While the former has been reported to affect perceived depth [19], the latter affects perceived position of the stimulus. Second, in addition to presenting our stimuli at high-contrast, we also presented them at lower contrast. This was motivated by the findings that MIPS magnitude scales with blurring of the stimulus and reduction in apparent contrast. We therefore expected larger perceptual displacements for low-contrast stimuli. Third, [12] reported results for early visual areas V1 to V3. In the current study we included several additional visual areas, including V3ab, hV4, LO, VO and TO (which comprises V5/hMT+; see [20]) to be able to study the effect on retinotopic representation across a wider range of areas. Fourth, potentially the difference in signal that comes with the shift in retinotopic representation is too small to be picked up by human neuroimaging methods. Yet advances in ultra-high field MRI [21] now allow for reliable recordings of human subjects at higher field strength which yield an improved signal-to-noise ratio. For the present study, we combined 7 Tesla [T] MRI with a custom-made surface coil optimized for high-resolution imaging of the human visual cortex [22] to obtain state-of-the-art recordings.

We recorded fMRI responses from early- and mid-level visual areas in humans while they were presented with pRF mapping stimuli that varied along two factors: the motion direction of the carrier pattern (3 levels: inward, outward and flicker motion) and the contrast of the mapping stimulus (2 levels: low and high stimulus contrast)(Figure 1). We hypothesized that carrier motion should displace population receptive field estimates against the direction of carrier motion. Furthermore, we expected greater and more reliable displacements for lower as compared to higher contrast.

## 2 Methods

### 2.1 Participants

Six healthy participants with corrected-to-normal vision were recruited for the study. All participants were students of Maastricht University. Recruitment was limited to participants who were MRI compatible and had been in an MRI scanner before to ensure high subject compliance. All participants were experienced in fixation tasks. The data for one participant (S1) were excluded from further analysis because mirror and projector settings in the scanner were updated between scanning sessions, rendering the data for different scanning sessions of this subjects incompatible in terms of eccentricity of the presented stimuli. Hence, five participants (4 females, 24-30 years old) were analyzed. All participants gave informed, written consent to participate in the experiment. The study was approved by the research ethics committee of the Faculty of Psychology and Neuroscience of Maastricht University.

**Figure 1:**
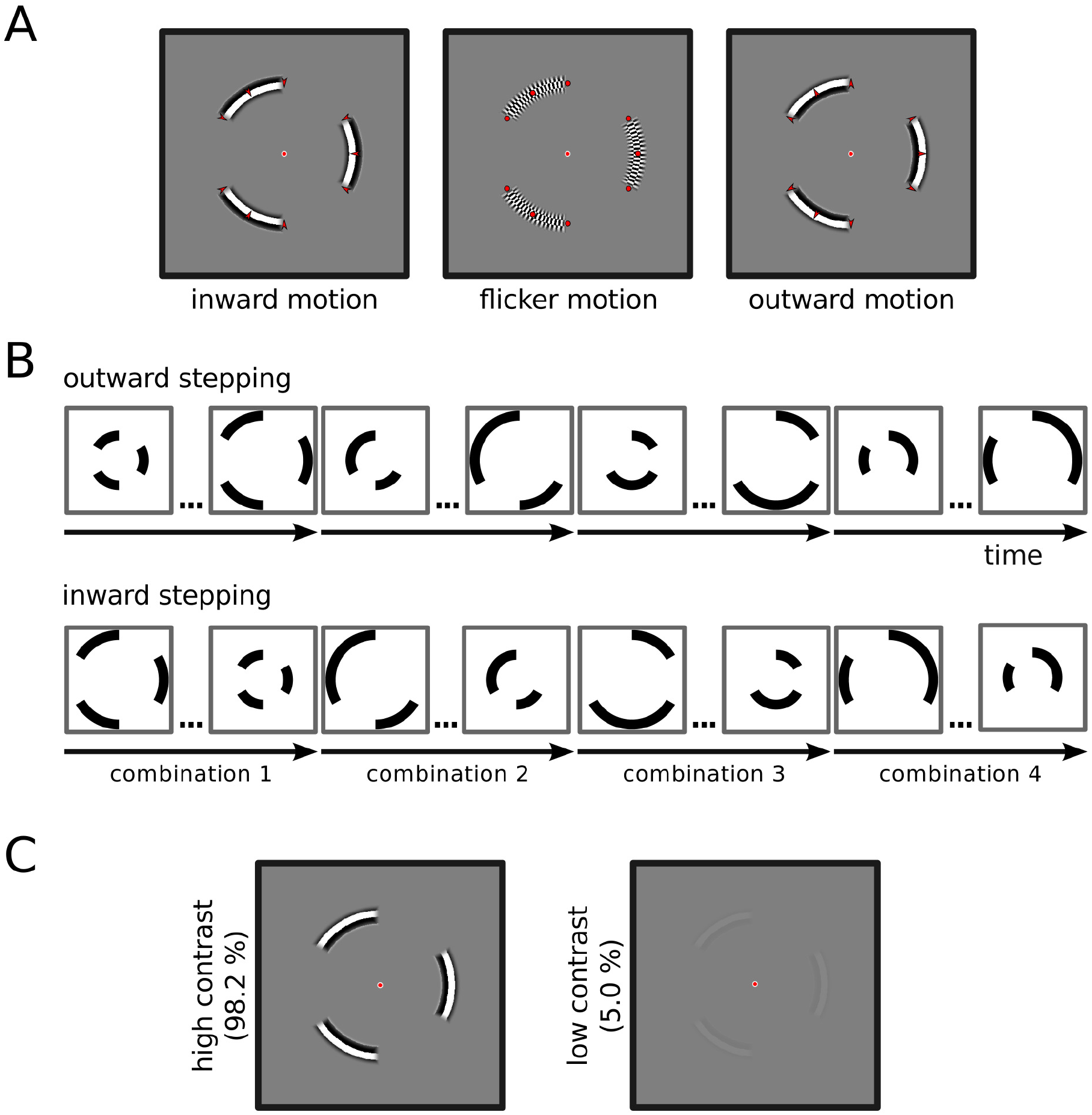
Overview of the experimental design. We systematically varied carrier pattern, aperture position and stimulus contrast. **(A)** Three types of carrier pattern. During inward and outward motion conditions, apertures revealed a radial square wave grating which drifted either inward or outward, while the aperture remained stationary. During flicker condition, apertures revealed a dartboard pattern, as classically used for pRF mapping. Red arrows and dots illustrate motion direction and were not shown during the experiment. **(B)** Aperture positions. Stimuli were presented behind stationary annulus sector apertures, indicated schematically as black sectors. Aperture position was systematically varied. There were four different annulus sector combinations (combination 1-4), covering different polar angles. These sector combination either stepped from low to high eccentricities (outward stepping) or from high to low eccentricities (inward stepping). Consecutive dots indicate in-between aperture position. **(C)** Stimulus contrast. All stimuli were presented at both high (98.2%) and low (5.0%) Michelson contrast.

### 2.2 Experimental design and stimuli

Visual stimuli were created and presented using the open-source software package PsychoPy (version 1.83.01) [23, 24]. Stimuli were projected on a frosted screen (using Panasonic projector PT-EZ570; Newark, NJ, USA; resolution 1920 × 1200; nominal refresh rate: 60 Hz). Subjects viewed the screen via a mirror attached to the surface coil. Button responses were registered using an MR compatible button box (Current Designs, 8-button response device, HHSC-2 × 4-C; Philadelphia, USA). Subjects participated either in two (S6) or three (S2, S3, S4, S5) scanning sessions of approximately 2 hours duration. Scanning sessions were on separate days. In session 01, we collected 4 runs of population receptive field mapping which was used to define regions of interest (ROI) as well as to derive area- and subject-specific hemodynamic response functions (HRF) (308 volumes each; independent ROI and HRF data). In sessions 02 and 03, we acquired 9 to 12 runs of pRF data for the main experiment at low and high luminance contrast (222 volumes each; low- and high-contrast pRF data). The different contrast levels were presented in separate sessions.

#### 2.2.1 Low- and high-contrast prf data

The pRF mapping stimuli consisted of annulus sector apertures presented at systematically varied positions. The choice of annulus sectors, as opposed to entire annuli [12], was motivated by our aim to manipulate perceived position, not perceived depth [19]. A single sector had a radial extent of 1.7°of visual angle and subtended 60 angular degrees such that the visual field was divided into six sectors in the angular direction, starting at the upper vertical meridian. At any moment, a combination of three sectors with equal eccentricity was shown. There were four possible sector combinations and each of the six sectors was shown during two of these combinations (see Figure 1B). Combinations were chosen to make the presentation of one sector maximally independent from presentation of other sectors. This was done by following a binary decision tree with four levels, picking the branches that led to exactly two successes and two failures. A binary decision tree with four levels has exactly six such branches and a single branch determined for one of the six sectors whether it was shown in a given combination (shown if success; not shown if failure).

For every combination, the eccentricity was systematically varied such that a combination started at a minimum radius of 3.4°and stepped up to a maximum of 8.5°, updating every 2 s (our TR) in 15 continuous steps. For the presentation of the first two and last two apertures the inner and outer edge, respectively, remained fixed. This resulted in an aperture step size of 0.34°. Combinations could either step from low to high eccentricities or, reversely, from high to low eccentricities and stepping direction was counter-balanced across functional runs. After completion of either 15 inward or outward steps, a different sector combination completed the same sequence of steps.

During inward and outward motion conditions, apertures revealed a radial square wave grating (spatial frequency: 1 cycle per degree) which drifted 5°of visual angle per s inward or outward, respectively, while the aperture remained stationary (Figure 1A). During the flicker condition, apertures revealed a dartboard pattern, as classically used for retinotopy and pRF mapping [17]. In radial direction, a dartboard element had an extent of 0.5°of visual angle, thus matching the spatial frequency of the square wave gratings during the inward and outward motion condition (1 cycle per degree). In the angular direction, dartboard elements subtended 3.36 angular degrees (360 / 107 = 3.36). This extent was chosen such that even at maximum eccentricity (8.5°of visual angle), the extent of the dartboard elements in angular direction was smaller than the extend in radial direction. This ensured that when the dartboard was flickered, elements were perceived to move in angular rather than radial direction. Elements were perceived to move either clockwise or counter-clockwise and perceived direction changed randomly over time [20]. The dartboard was flickered at a frequency of 5 on-off cycles per s such that the frequency was matched to the inward/outward motion conditions.

Gratings were presented either at 5% (low-contrast condition) or at 98.2% (high-contrast condition) Michelson contrast, which was the highest possible contrast in the scanner environment. In order to reduce increased responses to the aperture edge [13, 15], contrast was ramped following a raised cosine over a distance of 0.6 deg from the aperture edge. Luminous intensity was determined by varying the Psychopy settings for color from the most negative setting ([−1, −1, −1], i.e. “black”) to the most positive setting ([1, 1, 1], i.e. “white”) in 17 linear steps and concurrently measuring luminance inside the scanner environment using a photometer. Every of the 17 measurements was repeated twice and a cubic polynomial function was fit to the mean across measurements. The resulting polynomical function allowed us to estimate the intensities, given the settings in Psychopy, and to calculate the Michelson contrast according to (*L*1 − *L*2)/(*L*1 + *L*2), where *L*1 and *L*2 refer to the luminance of “white” and “black” segments, respectively.

To avoid adaptation to motion direction [10, 25], all stimulus presentations in the main experiment were restricted to brief periods of 400 ms. During the first and last 50 ms of the 400 ms presentation period contrast was ramped on and off following a raised cosine in order to reduce on- and offset effects. This was followed by 600 ms blank screen. During a TR (2 s), this sequence was shown twice. Every functional run consisted of 222 volumes and was divided into 3 parts (74 volumes). Each part showed one of the 3 conditions (outward drifting, inward drifting, flicker motion) and consisted of 7 volumes blank screen in the beginning and end as well as a sequence of 15 steps (either inward or outward stepping) for each of the four sector combinations (4 × 15 volumes = 60 volumes).

Participants were asked to fixate on a centrally presented dot throughout the entire experimental run. The fixation dot consisted of a red circle of 0.125°radius. To increase visibility of the fixation dot, it was surrounded by a white annulus of 0.19°radius [18]. To ensure that participants were complying with the fixation task, occasionally the central fixation dot changed color for 0.3 s to perceptually either darker or brighter shades of red. Participants were asked to indicate the direction of the change by pressing one of two buttons (button 1 for “darker”, button 2 for “brighter”).

#### 2.2.2 Independent ROI and HRF data

In an additional session, we obtained data to estimate population receptive field parameters independent of the main experiment. These data were used to define visual field maps, to select voxels for the main experiment and to determine area- and subject-specific HRF. We again systematically varied position and shape of the stimulus aperture, this time using three types of apertures: vertical bars, horizontal bars and circular sectors (“wedges”). All apertures were limited to a circular region of the display, 24°of visual angle in diameter, and presented against a mean luminance gray background. Vertical or horizontal bars traversed the visual field in 31 discrete steps. Vertical/horizontal bars stepped either from the left/top to right/bottom and back, or in reverse order. Both horizontal and vertical bars subtended 3°of visual angle in the stepping directions. In the direction orthogonal to stepping they were limited by the global circular aperture. The wedge aperture (angle width 45°, radius 12°of visual angle) completed a full circle in 32 steps, moving either clock-wise then counter-clockwise, or in reverse order.

The carrier pattern for all aperture types consisted of a dynamic, high-contrast pseudo-checkerboard as described by [26]. In comparison to traditional flickering checkerboard patterns, such a carrier pattern of concentric shapes varies in spatial frequency and phase over time and ensures stimulation of a broad range of visually responsive neurons [27]. The carrier pattern was always shown at 98.2% Michelson contrast (i.e., the highest possible contrast in the scanner environment). For every aperture type, contrast was ramped following a raised cosine over a range of 0.3°from the aperture border.

Aperture positions were updated with every TR (2 s). To avoid the appearance of apertures “jumping” from one position to the next, the stimulus was shown for 1.5 s only, followed by 500 ms blank screen. During the first and last 100 ms of the 1.5 s stimulation, contrast faded in or out following a raised cosine. Every functional run started and ended with 14 s blank screen. During every run, all three aperture types were shown. Vertical and horizontal bars each traversed the visual field twice; the wedge aperture traversed the visual field four times. The order of aperture types and traversing directions was counter-balanced across runs according to a balanced Latin square. Every two traversals of the visual field (back and forth) by one of the aperture types was followed by a period of 28 s blank screen.

As in the main experiment, participants were asked to maintain central fixation. Make-up of the fixation dot was identical to the one in the main experiment. The fixation dot occasionally changed color from red to yellow for 0.3 s and participants indicated a color change by pressing a button. When the aperture stimuli passed behind the fixation point, they spared out a circular region of 0.4°radius that revealed the fixation dot and part of the gray mean-luminance background. This was to avoid eye motion tracking the moving stimulus aperture [18]. To furthermore facilitate central fixation, an extended grid of thin lines was presented throughout the experimental run. The grid consisted of two diagonal, orthogonal lines crossing behind the fixation dot as well as circles with radii of 1°, 2.5°, 5°, 10°and 15°of visual angle centered on the fixation dot. Such a stable grid stabilizes fixation [28]. This grid was not shown in the main experiment to prevent participants from using it as a reference for location of the apertures.

### 2.3 Psychophysics experiment

To determine the perceptual displacement of the stimuli used in the fMRI experiment, all participants took part in a separate psychophysics session. This session followed a similar procedure as described by [18], with modifications where necessary due to differing stimuli. Like the fMRI experiment, all psychophysics stimuli were generated in PsychoPy (version 1.83.01) [23, 24]. Luminance measurements were repeated in the environment of the psychophysics lab, the polynomial function was refit and stimulus settings were adjusted to match the contrast levels employed in the fMRI experiment (5% and 98.2%).

Stimuli in the psychophysics session consisted of a radial sector aperture which subtended 120 angular degrees and had an extent of 1.7°. The aperture was centered on the upper vertical meridian at a distance of 5.95°from fixation, which represented the center eccentricity during the fMRI experiment. The carrier patterns were the same as in the fMRI experiment (outward, inward or flicker motion) and stimuli were again presented at two contrast levels (5% and 98.2% Michelson contrast).

In order to derive psychometric functions, we used the method of constant stimuli and a two-interval forced choice paradigm. On a given trial we presented two stimuli in temporal succession and then systematically varied the vertical distance between them. There was either no offset or the second aperture was 0.05°, 0.1°, 0.2°, 0.4°, or 0.8°above or below the first one. Stimuli were shown for the same duration as in the fMRI experiment (400 ms, of which contrast was ramped during the first and last 50 ms) and an interval of 500 ms blank screen was inserted between two subsequently shown stimuli. Subjects were asked to judge whether they perceived the aperture that was presented first or second as higher and communicated their judgment via button presses.

There were five possible combinations of subsequently presented stimulus pairs: outward then flicker motion, flicker then outward motion, inward then flicker motion, flicker then inward motion, flicker then flicker motion. The flicker then flicker combination was shown twice. An experiment block contained all vertical offsets crossed with all carrier pattern pairs, resulting in 11 (offset conditions) × 6 (motion combinations) = 66 trials per block. The order of trials was randomized. The two contrast-levels were presented in separate blocks. Each subject completed 8-10 blocks per contrast level.

### 2.4 Psychophysics analysis

Combinations that contained the same relative carrier patterns during the presentation of the two stimuli were combined. This reduced the number of combinations to three: flicker-flicker, outward-flicker and inward-flicker. For every vertical offset, we divided the number of times the subject judged the second aperture as higher by the total number of presentations at this offset. This yielded the proportion of perceiving the second aperture as higher which constituted our dependent measure. We then fitted a cumulative Gaussian psychometric function to these proportions using Bayesian inference based on samples from the posterior distribution as implemented in the Python version of the psignifit toolbox [29]. We used psignifit’s default priors for the ‘YesNo’ option, which reflect the belief that the point of subjective equality (PSE) falls within the range of measured data and, with decreasing probability, up to half the range above or below the measured data [29]. The PSE in our setting corresponds to the point where subjects perceived the two consecutively presented stimuli to be at equal height.

Based on individual psychometric functions, we determined the PSE for every carrier motion combination and subject. These individually fitted PSEs were used for inferential statistical analyses. We used the scaled PSE that is corrected for “guessing” and “lapse” rates estimated for every individual psychometric function. We averaged the scaled PSEs across subjects and calculated the standard error of the mean and 95% confidence interval.

### 2.5 MRI acquisition

Data acquisition was performed on a whole-body Magnetom scanner (nominal field strength 7 T, Siemens Medical Systems, Erlangen, Germany) at the Maastricht Brain Imaging Centre, The Netherlands. All images were acquired using a surface coil (4 channel transmit, 16 channel receive) custom-made for high resolution fMRI of the human visual cortex [22]. Images with anatomical contrast were acquired using the 3D MP2RAGE sequence (TR = 5000 ms; TI1/TI2 = 900/2750 ms; TE = 2.55 ms; FA1/FA2 = 5°/3°; FOV = 230 × 230 mm2; matrix size = 384 × 384; slices = 240; voxel resolution = 0.6 mm isotropic; GRAPPA factor = 2, partial Fourier = 6/8; phase encoding direction anterior - posterior) [30]. T2*-weighted functional data were acquired using a 2D gradient echo (GE) echo planar image (EPI) sequence (TE/TR = 20/2000 ms; in-plane FoV 144 × 144 mm^2^; matrix size 120 × 120; slices = 35; voxel resolution = 1.2 mm isotropic; nominal FA = 82°; echo spacing = 0.78 ms; GRAPPA factor = 2, partial Fourier = 6/8; phase encoding direction head - foot) using a protocol modified from [31, 32, 33].

### 2.6 Structural data preparation

Structural images were processed using FSL FAST [34], SPM’s bias correction [35], ITK-SNAP [36], FSL BET [37] and morphological operations [38]. Where not specified otherwise, default settings were used.

Since structural images were recorded using the visual surface coil, they had high signal close to the coil elements in the posterior part of the brain and gradually lower signal at anterior positions. We observed that this inhomogeneity affected the quality of the T1 map obtained from the MP2RAGE sequence: We found good quality in posterior parts and increasingly noisy responses and signal drop-out in anterior parts. Since we were only interested in early- and mid-level visual areas up to visual field map TO, and since noisy responses can negatively affect automatic segmentation algorithms, we first generated a mask to exclude voxels outside the occipital lobe.

We created the occiptal lobe mask by performing anatomical bias correction on the second inversion image and inputting the resulting image to FSL BET to obtain a first tentative brain mask. This mask often included the most posterior two thirds of the brain, where no signal drop-out was observed. We then used ITK-SNAP’s adaptive paint brush mode in combination with a graphics tablet (Intuos Art; Wacom Co. Ltd; Kazo, Saitama, Japan) to restrict this mask to the occipital lobe. An opening morphological operation (erosion followed by dilation) was applied to the mask after manual correction in order to remove high-frequency variations between slices that result from manual, slice-by-slice corrections.

The occipital lobe mask was applied to the T1 image and the masked T1 image was input to FSL FAST to obtain a white matter (WM) definition. This WM definition was used for boundary-based registration between functional and structural images (see below). Using the scalpel tool in ITK-SNAP, we divided the WM definition into two separate definitions for the left and right hemisphere. We dilated these WM definitions by one voxel and used the dilated definitions for surface reconstruction In BrainVoyager. Since voxel size was 1.2 mm isotropic and occiptal gray matter (GM) is 2-3 mm thick, this resulted in surface reconstructions along the middle of GM. The surface meshes were smoothed using BrainVoyager’s advanced mesh smoothing tools (restricting vertex displacement to 0.5) and inflated (500 fold). The resulting surfaces were used to display polar angle estimates obtained from the independant ROI and HRF data in order to delineate visual field maps.

### 2.7 Functional data preparation

Functional data were processed using BrainVoyager 20.0, SPM 12 [39], FSL 5.0 [40] as well as custom code in Python 2.7 [41, 38, 42] and in Matlab R2014a (The Mathworks Inc.; Natick, MA, USA). Where not specified otherwise, default settings were used. Functional images were pre-processed using the following operations: slice scan time and motion correction (SPM), high-pass filtering (FSL) and distortion correction (FSL topup). An overview of the pre-processing steps is provided by Figure 6.

For motion and distortion correction, we deviated from default settings. Functional images from each scanning session were motion-corrected using SPM 12 in three steps. First, the first image of runs in a given session was realigned to the first image of the first run of that session. Second, the images within each run were aligned to the first image of the run. Third, to avoid local minima, after these first two steps, a mean of all images in a session was calculated and images of that session were realigned to this mean. Motion correction was limited to voxels inside the brain based on an intensity-thresholded and manually corrected brain mask of the functional images. Note that the results of the three steps were combined into a single transformation that was applied to functional images to minimize interpolation artefacts.

EPI distortions of the functional images were corrected using FSL topup [43, 44]. The pairs of opposite phase encoding images acquired at the beginning of the first session of the main experiment (second scanning session) were input to topup to estimate the susceptibility-induced off-resonance field. The estimated field was then used to correct the distortions for all functional images of all scanning sessions.

To align the functional images acquired in different scanning sessions to each other, we calculated a mean image across time, separately for every scanning session (before EPI-distortion correction). We then ran an affine registration between the mean image of the second scanning session and each of the mean images of the other sessions using FSL FLIRT [45, 46] with 12 degrees of freedom. The resulting linear transformation was applied to all images recorded outside the second scanning session. We visually inspected the quality of the resulting alignment between the first run in the second session and all runs in the other sessions. Only then EPI distortion correction was applied to the transformed images, using the off-resonance field estimated in session 2. We applied a mask to all images, excluding voxels that had a mean intensity value below 30 in at least one of the sessions. This excluded voxels at the fringes of the slabs (due to minimal differences in slab placement across the three sessions).

Each run for the low- and high-contrast pRF data was split in 3 parts corresponding to the presented motion conditions. Since during each run (222 volumes) each of the 3 motion conditions were presented, this resulted in segments of 222 / 3 = 74 volumes. The division point between two motion conditions fell in the middle of a 14 volumes rest block such that carry-over effects from one motion condition to the next were minimal. All time courses were demeaned and averaged across runs. The low- and high-contrast pRF data were averaged separately for every motion condition.

To co-register functional data with the T1 image, we obtained an initial transformation estimate using the scanner’s positional information and fine-tuned this estimate using boundary-based registration [47] as implemented in BrainVoyager. The result was visually inspected for each subject by overlaying functional and anatomical images. Co-registration was implemented using nearest neighbor interpolation.

### 2.8 pRF modelling

#### 2.8.1 Overview

The pRF mapping data were analyzed as described in [17, 48, 49, 50, 51]. In brief, voxel time courses were assumed to be generated by an isotropic 2D Gaussian in visual space. Model responses were calculated by transforming the presented pRF stimuli to binary apertures (indicating presence or absence of a stimulus in space) and spatially convolving the apertures with an instance of the 2D Gaussian model. The Gaussian model included parameters for its position (*μ*_*x*_, *μ*_*y*_) and its size (*σ*; standard deviation of the 2D Gaussian) in visual space and the predicted model time course varied with assumed parameters [17]. In what follows, an exact description of the employed models is provided.

#### 2.8.2 Employed models

A voxel time course *y(t)* was modelled using the predicted fMRI time course *p(t)* of a given pRF model as predictor in a linear regression model according to

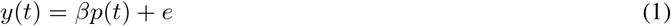

where p(t) is the predicted fMRI response for *t* = 1, …, *n* with *n* being the number of acquired volumes, *β* is a scaling factor that accounts for the unknown units of the fMRI signal, and *e* is measurement noise [17]. We solved for *β* using the ordinary least squares method.

The predicted fMRI time course *p(t)* for a pRF model was obtained according to

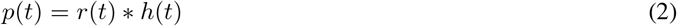

where r(t) is the time course response for a given pRF model. This time course was convolved with an HRF h(t) to account for the hemodynamic delay in the fMRI response. The employed HRF was either the canonical double-gamma function [52] or estimated subject- and area-specific functions (see below). The time course response of a pRF model was calculated as linear multiplication between the stimulus and the model pRF

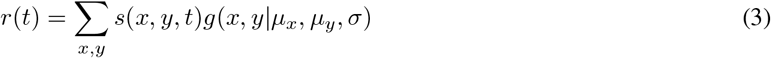

where s(x,y,t) describes the effective stimuli presented over time t and x and y represent positions in the visual field.

The pRF was modelled as an isotropic 2D Gaussian following

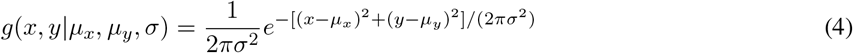

where *μ*_*x*_ is the preferred x-position, *μ*_*y*_ is the preferred y-position, *σ* is the size of the pRF. The parameters *μ*_*x*_, *μ*_*y*_, *σ* were determined for every voxel by minimizing the sum of squared errors (SSE) between the predicted and the observed fMRI time course [17]. Note that SSE was calculated on the same data as used for optimizing the parameters.

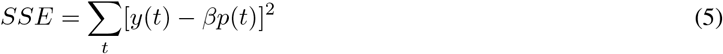

We obtained an approximation of percent signal change (% signal change) for every voxel by determining the maximum of the predicted time course and dividing by the mean signal time course over time.

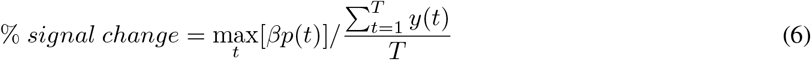

We also employed established extensions of the standard pRF model [49, 50]. These extensions included a compressive non-linearity as described by [50, 51]. For this extension the predicted fMRI time course was calculated as

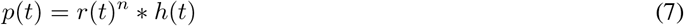

Raising the response element-wise to the power of *n* will act as a compressive nonlinearity if *n* < 1. Another extension was to add a suppressive surround using a difference-of-Gaussians (DoG) model as described by [49]. For this extension a voxel time course was modelled as

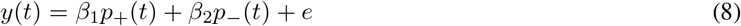

where *p*_+_(*t*) describes the response of the excitatory center and *p*_−_(*t*) the suppressive surround. We required that (i) *β*_1_ > *β*_2_, (ii) *β*_1_ > 0 and (ii) *β*_2_ < 0. Since we did not expect that all voxels show a center-surround organization, we used the classical pRF model for all voxels in which the surround model did not yield a lower SSE (equation 5). Subtraction of the surround Gaussian changes the effective size of the excitatory center. Therefore, instead of reporting *σ* for the pRF size we reported the half width at half maximum (HWHM). For voxels with the classical model, the HWHD maximum could be easily calculated as 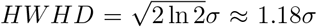. For voxels with a suppressive surround, HWHD was approximated empirically. The size of the suppressive surround was calculated as the distance between the point where the pRF reaches its minimum amplitude and the pRF center [49].

#### 2.8.3 Optimisation

Parameters of the pRF model were optimized in a two-step procedure. In an initial step, we created candidate model time courses on a grid and minimized SSE in a brute force search across candidate models (see below for specific parameter choices of the grid). Exhaustively searching a dense grid was achieved in a timely manner using multi threading and optimized Cython code. In the second step, we used non-linear optimization (Nelder-Mead algorithm) with constraints as implemented in the Python package lmfit [38, 53]. Using this strategy of combining an initial grid search with non-linear optimization mitigates the problem of finding *local* optima.

### 2.9 Analysis of independent ROI and HRF data

The independent ROI and HRF data served three purposes. First, we used them to obtain independent pRF estimates which were used to delineate early- and mid-level visual field maps. Second, we used them to derive subject- and field-map specific HRF. Finally, the data were used for voxel selection in the main experiment.

#### 2.9.1 Visual field map delineation

For this analysis only, time courses were not averaged across runs and we applied Gaussian temporal (kernel width = 2 s) and spatial (kernel width = 1.2 mm) smoothing to the pre-processed EPI images. The spatial smoothing was motivated by the prior assumption that pRF parameters vary smoothly over space, which aids the process of manually delineating visual field maps.

To delineate visual field maps, we estimated a voxel-wise pRF model with a compressive non-linearity. The code implementing this estimation is publicly available [54]. Candidate pRF model time courses were created on the following grid: 64 radial position (0°-12°eccentricity) × 64 polar positions (0-2*π*) × 40 pRF sizes (0.2°-16°) × 5 exponents (0.1, 0.2, 0.4, 0.7, 1.0). The resulting pRF parameters were projected onto the inflated, reconstructed hemispheres. This allowed us to delineate visual field maps on every hemisphere by manual drawing following established strategies. In particular, visual field maps V1, V2v, V2d, V3v, V3d were clearly described by reversals in polar angle. A combined map of V3a/b was delineated using criteria described in [55, 56] and no effort was made to obtain separate maps for V3a and V3b. To delineate hV4 as well as VO1 and VO2, we followed landmarks and strategies described in [57]. Field maps LO1, LO2, TO1 and TO2 were delineated following procedures described in [20]. Additionally, we observed that visual field maps TO1 and TO2 were often marked by a substantial drop in the exponent for the compressive nonlinearity. To increase the number of participating data points, we collapsed the identified visual field maps into V1, V2, V3, hV4 [58], V3ab, LO, VO and TO. Resulting delineations, along with the pRF properties that guided the delineations, are shown for one representative hemisphere in Figure 7. The combined field maps were projected back into volume space (−1mm to +1mm from the reconstructed surface) and voxels for a common field map were grouped across the left and right hemisphere.

#### 2.9.2 Hrf estimation

To obtain subject- and field-map specific HR functions, we proceeded in two steps [48, 59, 18]. First, we estimated the pRF parameters using the canonical double-gamma function [52] as the HRF. For this estimation, we employed the difference-of-Gaussians model with suppressive surround and a brute force search for optimization (64 radial position [0°-12°eccentricity] × 24 polar positions [0-2*π*] × 40 pRF sizes [0.2°-16°] × 15 surround ratios [1.1, 1.2, 1.3, 1.4, 1.5, 1.6, 1.8, 2.0, 2.2, 2.4, 2.6, 2.9, 3.2, 3.5, 4.0], where a ratio indicates the size of the surround compared to the size of the center). For every visual field map we then selected voxels that had a coefficient of determination (R2) of at least 0.4 on the training data. This was done to exclude visually unresponsive voxels from the HRF estimation. In case a visual field map had more than 500 voxels with R2 greater than 0.4, the 500 voxels with the highest R2 value were selected to keep the computing time practicable.

Second, for selected voxels we kept the estimated pRF parameters fixed and varied the parameters of a double-gamma function with the objective of minimizing the SSE between the predicted and observed voxel responses. Hrf parameters were optimized using a brute force search on a grid of HRF parameters (5 grid positions for the delay of the peak [5.0 - 7.0 s], 10 positions for the delay of the undershoot [16.0 - 20.0 s], 5 positions for dispersion of the peak [0.1 - 1.6], 5 positions for dispersion of the undershoot [0.1 - 1.6] and 16 positions for the peak to undershoot ratio [5.0 to 1.0]). The code implementing this optimization is publicly available as part of the HrfOpt package [60]. To obtain a single field map-specific HRF, we averaged the optimized HRF parameters across all voxels in the field map that were included in the analysis (maximally 500 voxels). The resulting HR parameters that were employed for the analysis of the low- and high-contrast pRF data are shown in Figure 8.

#### 2.9.3 Voxel selection

We were interested in selecting only voxels that were visually responsive and, in particular, responsive to the eccentricities employed in the main experiment. We implemented these desiderata by performing a pRF analysis with compressive non-linearity on the independent ROI and HRF data to determine *μ*_*x*_,*μ*_*y*_,*σ* and *n* for every voxel. We then only selected voxels for the main experiment that had an R2 value of at least 0.4 which served as an indicator of “visual responsiveness”. Based on the pRF estimates we also calculated the expected maximal response *r* to the stimuli s(x,y,t) presented in the main experiment according to:

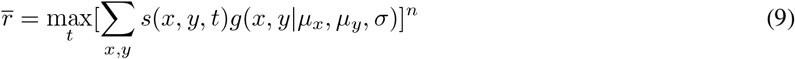

Expected values could range from 0 (unresponsive) to 1 (fully responsive). We only considered voxels with a predicted response of at least 0.5. Finally, we required all voxels to be located in one of the delineated visual field maps.

Three criteria thus determined the selection of voxels for the main experiment: (i) voxels needed to have an R2 value of at least 0.4 on the independently acquired pRF data, (ii) voxels needed to have an expected response to the stimuli in the main experiment of at least 0.5, and (iii) voxels needed to be located in one of the field maps.

### 2.10 Analysis of low- and high-contrast pRF data

We estimated a voxel-wise pRF model with suppressive surround separately for every carrier motion (inward, outward, flicker) and for the low- and high-contrast data. The code implementing this estimation is publicly available as part of the PyPrf feature package [61]. We constructed candidate pRF model time courses on a grid (64 radial position [0°-12°eccentricity] × 24 polar positions [0-2*π*] × 40 pRF sizes [0.2°-16°] × 15 surround ratios [1.1, 1.2, 1.3, 1.4, 1.5, 1.6, 1.8, 2.0, 2.2, 2.4, 2.6, 2.9, 3.2, 3.5, 4.0]) and minimized SSE in a brute force search. Subsequently, we used non-linear optimization to further adjust the *μ*_*x*_, *μ*_*y*_ and *σ* parameters. The ratio of size-of-sorround to size-of-center was carried over from the brute force search and kept fixed during non-linear optimization. During both brute-force and non-linear optimization the center of the estimated pRF was constrained to be in the visually stimulated part of the visual field. During all analyses for the main experiment we employed the subject- and area-specific HR functions derived above. In a second, independent analysis we repeated the analysis outlined above but this time without the suppressive surround option.

To compare pRF parameters across motion conditions, voxels in each field map were pooled across subjects. Results per field map were summarized using the median as a measure of central tendency and 68% confidence intervals were obtained by 50,000 fold bootstrapping of medians to summarize variability across voxels.

### 2.11 Simulation of shifted population position codes

To investigate the potential effect of changes in pRF position on perceptual position shifts, we implemented a model-based decoding analysis. For this analysis we grouped the pRF parameters found in the independent pRF mapping experiment across all subjects and then applied systematic shifts to the pRF centers in the order that was observed empirically: [0.1, 0.1, 0.1, 0.3, 0.2, 0.6, 0.17, 0.0] across V1, V2, V3, V3a/b, hV4, LO, VO, TO to simulate inward motion and [−0.1, −0.1, −0.1, −0.3, −0.2, −0.6, −0.17, −0.0] to simulate outward motion. To simulate flicker motion, no shifts were applied. We used the resulting pRF to simulate the responses 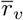 of each voxel that would be produced by the different stimuli shown during the psychophysics experiment. The simulated response patterns were projected into the visual field by calculating the 2D projection *M* as

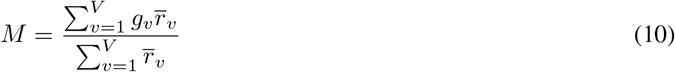

where *g*_*v*_ = *g*(*x, y µ_x_, µ_y_, σ*) is the 2D Gaussian for a pRF with *μ*_*x*_, *μ*_*y*_, and *σ*), 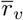 is the simulated response and *V* is the total number of voxels in a given field map. *g*_*v*_ was assumed to be fixed across different motion conditions, reflecting the assumption that retinotopy is not flexible [12]. Instead 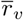 varied across flicker, inward and outward motion as described above.

This resulted in one visual field projection *M* per psychophysics stimulus, motion condition and field map. For every projection we calculated the expected eccentricity value *E*(*Ecc*) (also referred to as the center of mass; [12]) according to

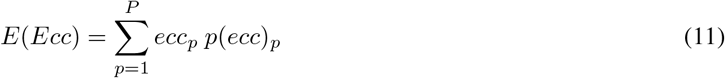

where *P* is the total number of pixels in a projection image, *ecc*_*p*_ is the eccentricity value at a given pixel *p*, and *p*(*ecc*)_*p*_ = *M*_*p*_ is the normalized value of a projection at a pixel *p* (normalized such that 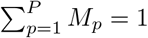).

For a given projection, we also calculated the variance of the expected eccentricity value according to

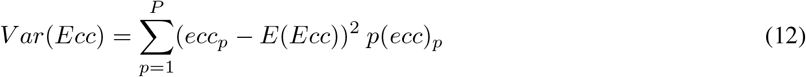

Intuitively, when we compare two projections *M*_1_ and *M*_2_ for two stimuli *s*_1_ and *s*_2_ then the stimulus for which the projection has a higher expected eccentricity value should be perceived as higher on a proportion of trials *p* > 0.5. This intuition was captured via a cumulative Gaussian function [62] and we calculated the proportion *p* of trials where the second stimulus is perceived as higher than the first according to

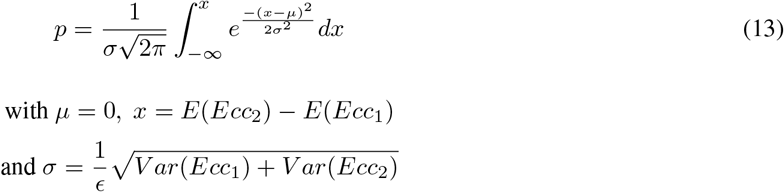

where *ϵ* is an efficiency parameter that was set to 0.1 [62]. This allowed us to calculate a proportion *p* for all the stimulus pairs shown in the psychophysics experiment. The same fitting as described in the psychophysics methods was used on these proportions to estimate the PSEs for every motion condition combination and visual field map.

In a second pass, we repeated the analysis above but this time, in addition to assuming shifts in pRF center, we also assumed differences in pRF size between motion conditions. In particular, we assumed an increase in pRF size by [0.08, 0.11, 0.26, 0.51, 0.66, 0.51, 0.61, 0.54] across V1, V2, V3, V3a/b, hV4, LO, VO, TO to simulate inward motion and a decrease of the same order to simulate outward motion. To simulate flicker motion, no changes were applied.

## 3 Results

### 3.1 How do motion and contrast influence perceived position?

Figures 2A and B show group-averaged psychometric functions for high- and low-contrast stimuli, respectively. For high-contrast stimuli, the fitted psychometric functions are very similar for the different carrier motion combinations and we do not observe a systematic shift in perceived position. By contrast, for low-contrast stimuli the fitted psychometric functions are clearly separated and the arrangement of the curves is consistent with previous studies that reported a shift of perceived position in the direction of motion [1, 2, 3, 4, 5, 6, 7, 9]. This pattern is also reflected in the individual psychometric functions for single-subject data (see Figure 9 and 10): While there is a systematic perceptual displacement in the direction of motion for low-contrast stimuli, no such pattern is evident for high-contrast stimuli.

**Figure 2:**
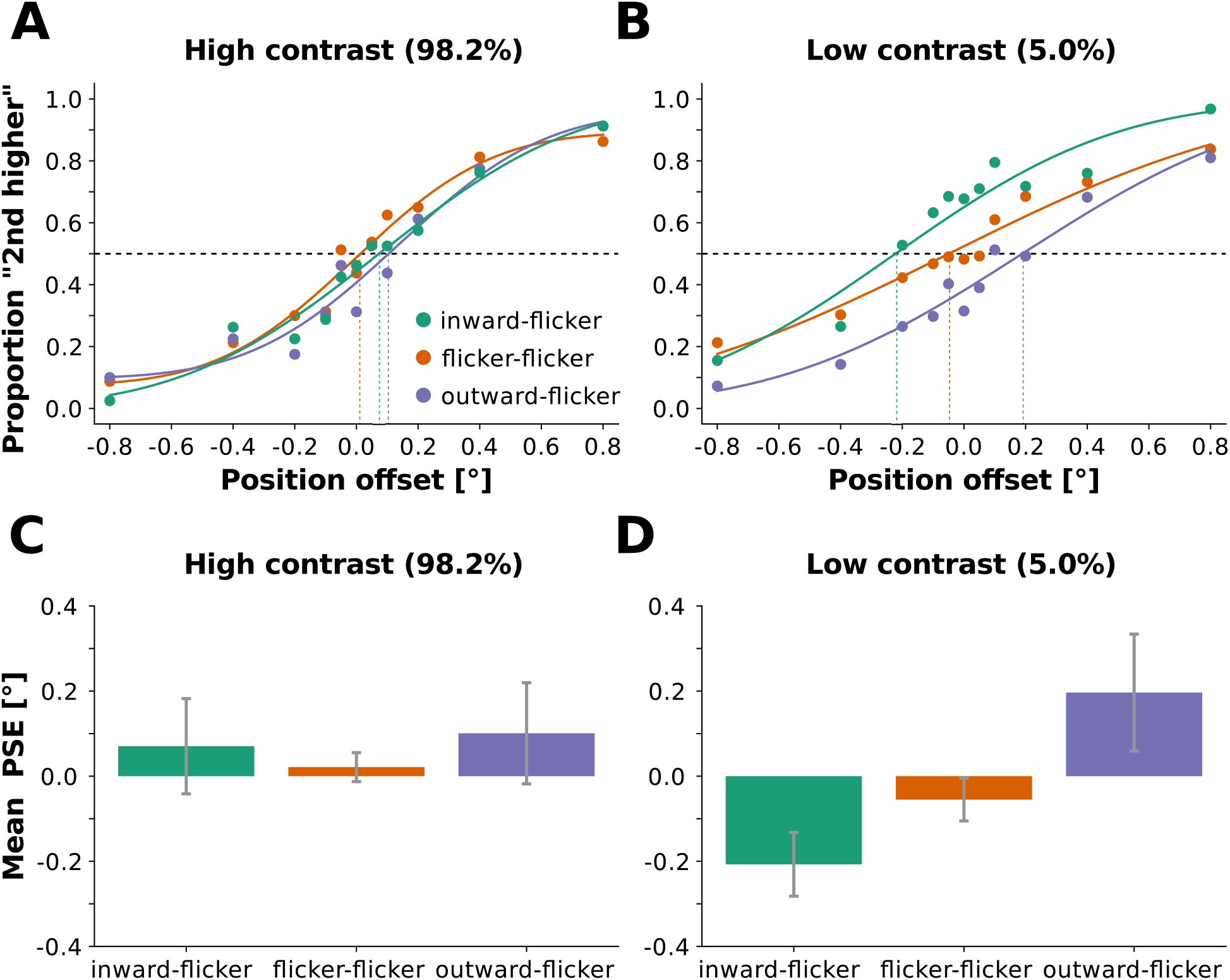
Displacement of perceived stimulus position in the direction of motion only observed for low-contrast stimuli. We varied the physical position offset between two subsequently presented stimuli and the carrier pattern of the stimulus (flicker, inward and outward motion). Participants indicated whether they perceived the first or second stimulus as higher. **(A)** Data and fitted psychometric functions for high-contrast stimuli averaged across participants. Dots represent data and curves represent fitted functions. The fitted curves represent responses for the sequence of inward then flicker motion (green), flicker then flicker motion (orange) and outward then flicker motion (violet). The psychometric functions largely overlap. **(B)** Data and fitted psychometric functions for low-contrast stimuli. Same conventions as in **(A)**. Responses for sequences of inward and flicker stimuli (green) were shifted upward, indicating that subjects more frequently perceived the second (flicker) stimulus as higher than the first (inward). By contrast, responses for sequences of outward and flicker stimuli (violet) were shifted downward, indicating that subjects more frequently perceived the second (flicker) stimulus as lower than the first (outward). Flicker-flicker stimulus sequences perceptually assumed an in-between position (orange). **(C)** Point of subjective equality (PSE) for high stimulus contrast. Bars represent average PSE across subjects. Error bars represent 95% confidence interval. We did not find systematic perceptual displacements. **(D)** PSE for low stimulus contrast. Same conventions as in **(C)**. We found no perceptual displacements for flicker-flicker combinations (orange). Outward motion displaced the perceived stimulus position in the direction of motion (violet), increasing the PSE. Reversely, inward motion led to a decrease in perceived position (green).

Figures 2C and D plot the group mean PSEs (and 95% confidence intervals) obtained from psychometric functions that were fitted to each participant and pattern combination individually for high- and low-contrast stimuli. The group mean PSEs confirm the trend that was already apparent from the group-averaged psychometric functions. In the low-contrast condition, mean PSE for the outward-flicker combination was 0.20° 0.14°and was significantly different from the veridical position difference (since the 95% CI does not include 0). Reversely, mean PSE for the inward-flicker combination was −0.21° 0.07°and also significantly different from 0. In the high-contrast condition, no mean PSE for any combination was significantly different from 0. Furthermore, the mean PSEs for the outward-flicker as well as the inward-flicker combination were both positive with 0.10° 0.12°and 0.07° 0.11°, respectively. We thus did not find the perceived position to be shifted in any systematic direction for the high-contrast data.

### 3.2 How do motion and contrast influence pRF properties?

We found that the effect of carrier motion on pRF properties differed depending on the stimulus contrast and visual field map (Figure 3). For high stimulus contrast, we found no substantial changes in eccentricity for areas V1 - V3, replicating the finding by [12]. For higher visual field maps, starting from V3ab, however, we observed higher preferred eccentricity values for outward compared to inward motion - with the highest difference, 0.7°, in field map TO. Flicker motion led to values intermediate to those obtained for inward and outward motion. For pRF sizes we observed a systematic decrease for outward as compared to inward motion across all visual field maps, with increasingly higher differences for higher visual field maps. Beyond V3, these effects were substantial with differences from 0.76°in V3a/b to 2.83°in TO. pRF size values for flicker motion were in-between values for inward and outward motion and often closer to outward motion. Percent signal change was decreased for outward compared to inward motion, which is consistent with a bias for motion towards fixation that has previously been reported for eccentricities below 14°in humans [13].

For low stimulus contrast, we found lower preferred eccentricity values for outward as compared to inward motion across all visual field maps, with the notable exception of field map TO. This corresponds to a shift of the pRF center in the direction opposite to motion, which is consistent with results from the animal literature [10, 16]. Eccentricity decreases were in the range from 0.20°to 1.23°and the decrease was very consistent across voxels as indicated by small error bars. As for high-contrast stimuli, we observed a systematic decrease of pRF sizes across all visual field maps for outward as compared to inward motion. These decreases were higher for higher visual areas and in the range from 0.15°to 1.53°. As before, the flicker motion condition usually assumed values in-between the inward and outward motion conditions. As for high-contrast stimuli, percent signal change was decreased for outward compared to inward motion. Percent signal change for flicker motion was much higher than for either inward or outward motion, especially in visual field maps V1, V2, and V3.

**Figure 3:**
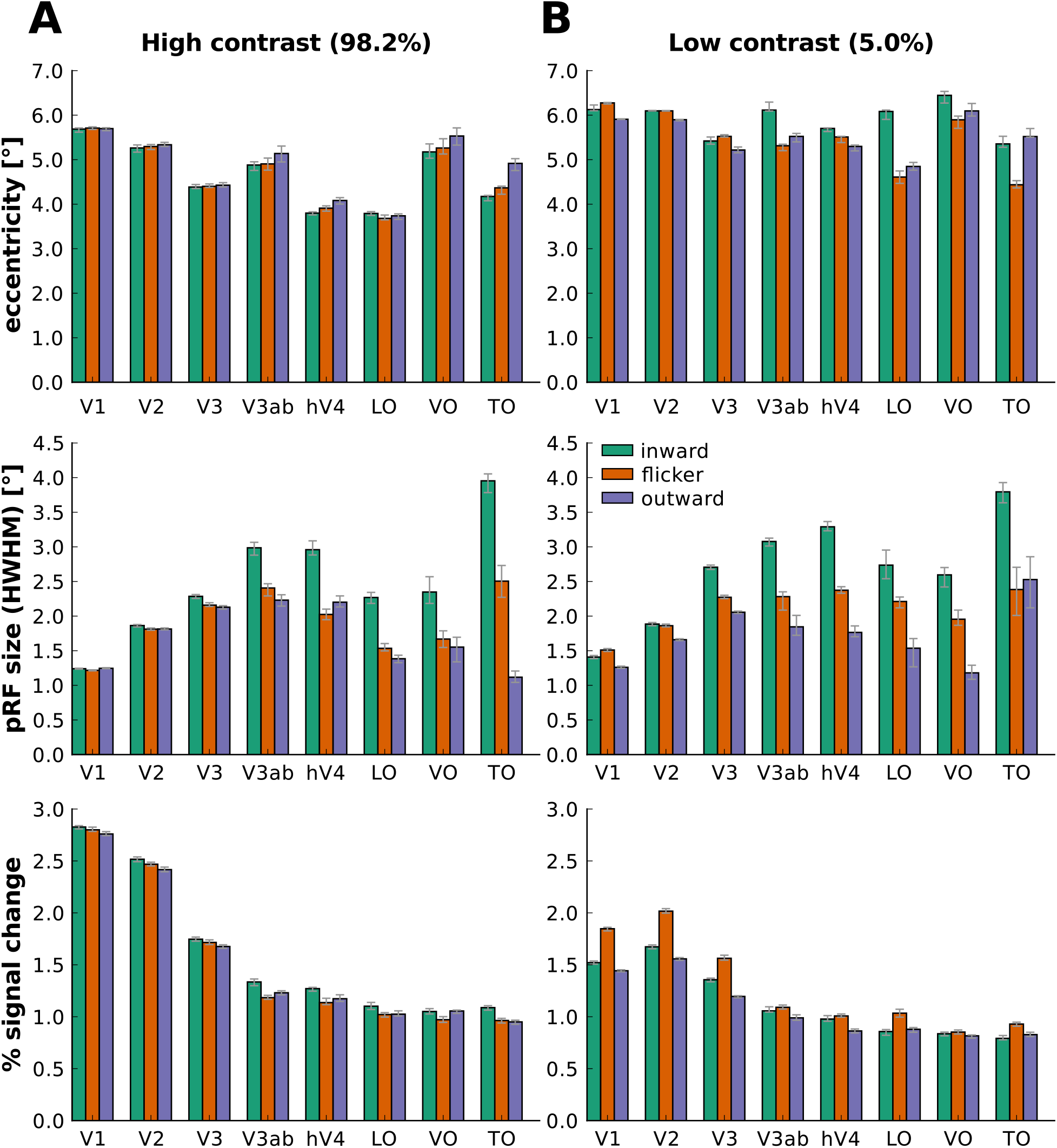
Effect of carrier motion on pRF properties depends on stimulus contrast and visual field map. Each bar represents a pRF property (median across voxels 68% confidence interval) in different visual field maps (V1 to TO) during one of three motion conditions: inward (green), flicker (orange) and outward (violet) motion. Panels **(A)** and **(B)** show results for high- and low-stimulus contrast. Different rows show different pRF properties: eccentricity of the pRF center (in °of visual angle), pRF size (HWHM, °of visual angle) and approximated % signal change.

The results described above were obtained for fitting the pRF model with the suppressive surround option. This allowed us to compare the effect of carrier motion also on suppressive surround size and the suppression index. Since the number of voxels that had a suppressive surround was low for visual field maps outside early visual cortex (less than 25 voxels for all subjects together), we only show the results for V1-V3. As shown in Figure 11, we observed larger suppression indices for outward as compared to inward motion. Furthermore, mirroring the observations for pRF size, we found lower sourround sizes for outward motion than for inward motion. Finally, when fitting the pRF model *without* suppressive surround, we obtained very similar results as before (Figure 12).

### 3.3 Can pRF center shifts account for perceptual shifts?

If the pRF center shifts in the direction opposite to motion, as they do for low-contrast stimuli, how can we account for the observation that, perceptually, the apertures appear displaced in the direction of motion? Figure 4 offers an explanation in form of a model for these seemingly contradictory findings. The model makes two essential assumptions: (i) during the MIPS, pRF centers in lower-level visual areas are shifted against the direction of motion; (ii) at the same time, retinotopy is *not* flexible in the sense that connections from lower- to higher-level visual areas are not dynamically updated during the 400 ms of stimulus presentation.

Figure 4A and B demonstrate how, given pRF shifts opposite to motion direction, lower-level visual areas send position codes that indicate a shift in the direction of motion. During natural viewing conditions, pRF centers are not displaced and lower-level visual voxels send position codes congruent with the physical position of a stimulus (middle column). However, during inward motion, pRFs will be shifted outward (left column) meaning that units that usually respond to stimuli at an eccentricity of, for example, 5°of visual angle (indicated by “orange” stimulation), will now respond to stimuli at an eccentricity of 5.2°visual angle (“green” stimulation). At the population level, the code that lower-level voxels of different eccentricities will send will signal a position of the stimulus that is slightly more foveal than the actual physical stimulus (see Figure 4B). During stimulation with outward motion, reverse conditions and predictions apply (see Figure 4A and B, right column).

**Figure 4:**
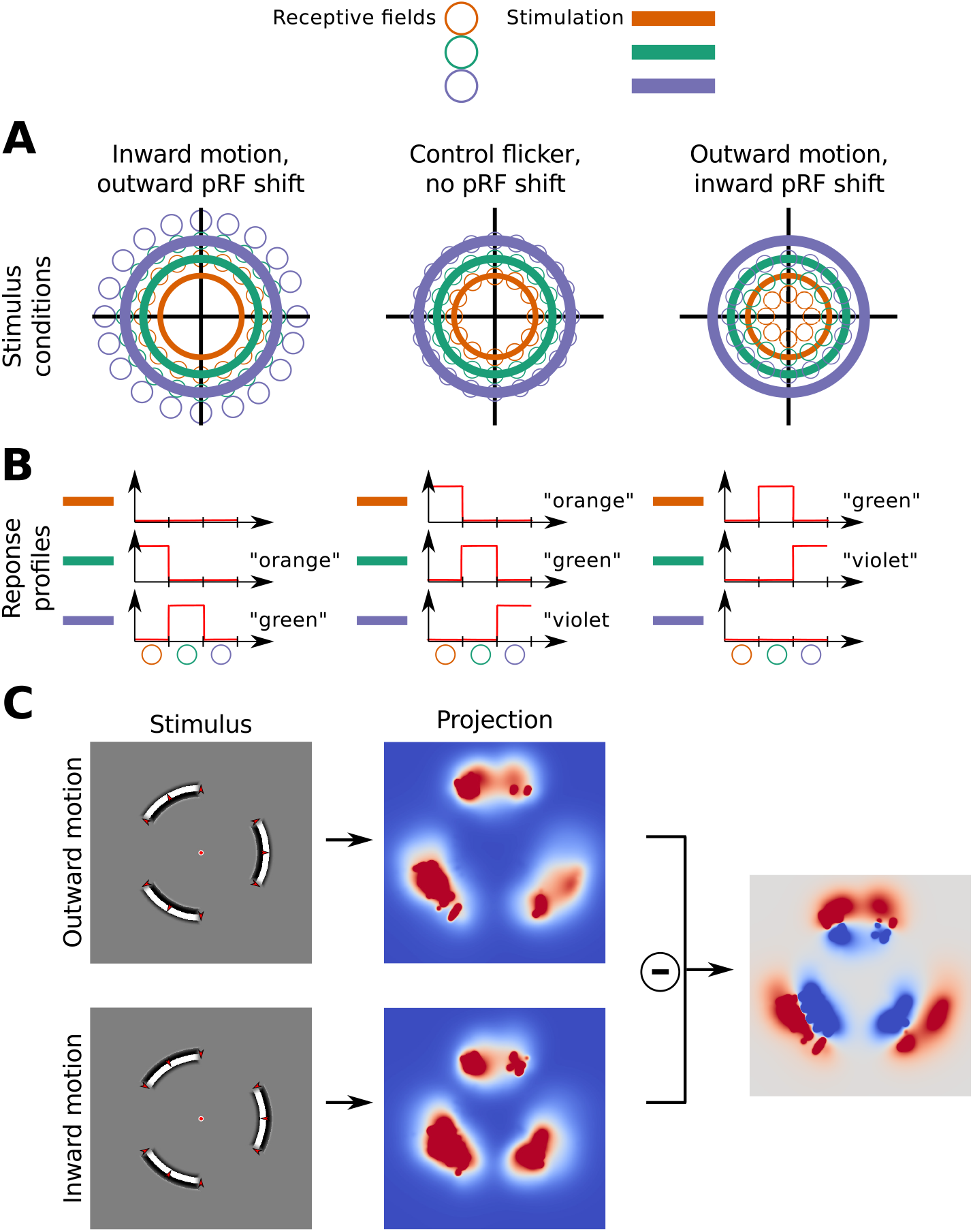
Simulated perceptual consequences of pRF shifts in the direction opposite to motion. **(A)** During natural viewing conditions, pRF centers are not displaced and lower-level visual neurons send position codes congruent with the actual position of a stimulus (middle column). However, if pRF centers are shifted outward during inward motion (left column), neurons that usually respond to stimuli at an eccentricity of, for example, 5°of visual angle (indicated by “orange” stimulation), will now respond to stimuli at an eccentricity of 5.2°visual angle (“green” stimulation). Reversely, if pRF centers are shifted inward during outward motion (right column), they will respond to more foveal stimulation than usual. **(B)** During inward (outward) motion neurons send a population level code that signals a position of the stimulus at slightly lower (higher) eccentricities than its actual position. For example, during peripheral (“violet”) stimulation, the neuron population will send a more foveal (“green”) code. **(C)** The shifted population level codes observed in **(B)** can be turned into quantifiable, perceptual predictions via visual field projections.

Figure 4C simulates the perceptual consequences of shifted population position codes. We assume the activation of pRF units is connected to (perceived) stimulus position via a visual field projection. The projection describes the mapping between the activation of a unit and the (perceived) presence of a stimulus at a certain position of the visual field. The mapping is learned during natural viewing conditions and is assumed not to change during inward (outward) motion, because connections between low-level and high-level neurons do not reorganise at the time scale of stimulus presentation. As a consequence, if during inward (outward) motion a population of units in a lower-level visual area signals a more foveal (peripheral) code than usual and connections between low- and high-level areas do not reorganise, then this will be interpreted (perceived) as the presence of a more foveal (peripheral) stimulus. This corresponds to a shift of perceived position in the direction of motion of the stimulus.

We performed a model-based decoding analysis to test whether the mechanism of shifted population position codes can reproduce perceptual shifts in the order that we observed empirically. In the low-contrast condition, we observed empirical pRF center shifts between inward and outward motion in the order of 0.20°to 1.23°across different field maps, while the perceptual shifts between inward and outward motion were in the order of 0.4°. Figure 13A shows the results of our simulation. We obtained perceptual shifts in the order of 0.1°to 0.5°depending on the visual field map. For early visual field maps V1-V3 the simulated perceptual shifts were too small to account for our empirical findings. However, the simulated perceptual shifts obtained in higher visual field maps approximate the perceptual shifts found empirically. We also investigated the effect of pRF size changes on perceptual shifts since we observed larger pRF sizes for inward compared to outward motion. Figure 13B shows that if we assume pRF size differences in addition to pRF center shifts, the simulated perceptual shift is enhanced. We discuss the potential contribution of pRF size differences to perceptual shifts below.

## 4 Discussion

We found that coherent motion signals impact both the perceived position of a stimulus as well as the estimated pRF position and identified stimulus contrast as an important mediating factor. Our study replicates findings from a previous fMRI study showing that preferred eccentricity does *not* change with carrier motion presented at high stimulus contrast in early visual areas V1 - V3. However, our study extends earlier findings in four important ways. First, we performed psychophysical measurements and only found MIPS when stimuli were presented at low-contrast but not at high-contrast. Second, we show that at low contrast, where perceptual displacements occur, pRF are shifted in the direction opposite to motion, which is consistent with findings from the animal literature [10]. Third, we study the effects of carrier motion on field maps beyond V3. Finally, we find effects of carrier motion on other pRF properties such as pRF size and suppressive surround size. We discuss these extensions in turn.

We observed substantial perceptual displacements only for low-contrast stimuli, not for high-contrast stimuli. Based on this finding, we propose contrast as an important factor that influences the magnitude of perceptual position shifts. This proposal can account for previous findings of larger MIPS with blurred stimulus edges [4, 9] or increased spatial blurring of the stimulus [4] since blurring decreases the effective stimulus contrast. Furthermore, this proposal is consistent with the object-tracking model that was suggested as a unifying framework for position and motion perception [9]. Accordingly, the human visual system implements an instance of an object-tracking system that attempts to infer the generative causes of retinal motion signals [9]. Lower stimulus contrast decreases the available sensory evidence and increases signal uncertainty. With increased uncertainty, the object-tracking system is more likely to falsely attribute retinal motion signals to object motion (resulting in the perceived position shift) instead of attributing it to the actual cause, pattern motion (i.e. the drifting square wave grating). Reversely, increased stimulus contrast reduces signal uncertainty, allowing the object-tracking system to correctly attribute retinal motion to pattern motion.

We found that when substantial perceptual displacements occurred, this was accompanied by pRF center shifts in the direction opposite to motion. Based on this finding and a corresponding simulation, we suggest that the perceptual shifts can be accounted for, neuronally, by shifted population position codes in early visual cortex. This result fills an important empirical gap between psychophysical, electro-physiology and fMRI research. Previously, it was unclear whether differences in findings between humans and cats/monkeys were due to differences in recording methods or inter-species differences in the underlying neuronal mechanism. We propose that the neuronal mechanism underlying MIPS is shared across human and non-human animals. The reason that this mechanism had previously not been discovered in humans was that perceptual shifts in response to high-contrast motion and their underlying neuronal mechanisms were probably too small to be measured using neuroimaging methods.

Our work suggests a synthesis of two different accounts that have been used to explain MIPS. Arnold et al. [7] have termed these two explanations the “RF account” and the “Contrast Modulation account”. According to the RF account, motion changes the region of retina to which a population of neurons responds. The Contrast Modulation account suggests that motion influences spatial coding via interactions that modulate apparent contrast [7]. These accounts were previously viewed as opposing each other but we propose that they should be considered connected. We demonstrate that perceptual displacements are linked to contrast (Contrast Modulation account). At the same time, we show that these perceptual displacements coincide with shifts of the pRF in the direction opposite to motion (RF account). This implies that any comprehensive account of motion and position coding in the human visual system should unite both explanations.

Two previous fMRI studies [63, 64] found that patterns of fMRI responses in early visual areas reflected physical stimulus position while higher areas like V3a, LO or hMT+ (included in TO) represented the perceived position. At face value, these findings are consistent with the pRF eccentricity changes that we observed for high-contrast stimuli. However, the previous studies used multivariate pattern analysis (MVPA). Therefore, it is difficult to clarify how exactly the results obtained in these studies and our study are related since several kinds of ambiguity are inherent to MVPA [65]. In particular, due to geometric ambiguity it is unclear whether dis/similarity of response patterns was based on overall activation alone or also orientation of the multivariate vectors. In addition, there is representational ambiguity which makes it impossible to determine whether position was the source of variation that was driving the observed degree of correlation between response patterns in those studies. By contrast, using a voxel-wise analysis, the pRF model allowed us to explicitly model position in visual space as the source of variation that was driving fMRI responses and avoided the ambiguities surrounding MVPA [65].

Several studies have found that the trailing edge of a motion stimulus evokes larger responses than its leading edge, at least for early visual cortex (V1-V3) [11, 12, 13, 15] but not for V5/hMT+ [13]. Could this trailing edge effect or “aperture-inward” bias account for some of the pRF property changes that we have observed here? We think that several considerations speak against a strong influence of the aperture-inward bias on our results.

First, it is unclear how the aperture-inward bias scales with the size of the aperture. We note that in our experiments, we employed apertures with an extent of 1.7°in the direction of motion, while studies that reported the aperture-inward bias used apertures that were considerably larger (4°-14°in [15] and 11°-22°in [13]). Several studies [13, 14, 15] have reported that responses decrease gradually with distance from the aperture edge. This decrease appeared to be approximately linear [14] at the employed aperture sizes. When employing smaller apertures, as we did here, this effect might scale to the new size of the aperture in which case a linear decrease from trailing to leading edge would be expected. However, it is equally possible that no decrease in responses is observed at all for small apertures since all receptive fields are in proximity to the aperture edge and the resulting difference between trailing and leading edge is minimal. This could be tested in future research by keeping one side of the aperture fixed and systematically increasing the extent of the aperture for different conditions.

Second, our stimulus settings and experimental design should have minimized aperture-inward bias. Moving dot stimuli seem to lead to larger aperture bias (in the order of 1.0 - 1.5 % signal change) [13, 15] than drifting grating stimuli (in the order 0.3 - 0.5 % signal change) [11, 12]. This can be attributed to the fact that moving dot stimuli unavoidably show a small asymmetry in dot onsets and offsets at the trailing and leading edge. Drifting grating stimuli, like we used in the current study, are unaffected by this problem and should show lower bias. Likewise, our experimental design and analysis should have reduced the bias. Directly contrasting the effects of opposing motion directions leads to larger effects than contrasting the effects of each motion direction with a mean-luminance background first and only then calculating the difference between the two effects [12]. This is because the hemodynamic nature of the fMRI signal acts like a compressive nonlinearity [12].

Third, all previous studies reporting aperture-inward bias presented only a single stimulus. It is therefore an open question how aperture bias affects pRF mapping experiments with varying stimulus apertures. To understand how the bias might affect the estimation of pRF properties, we ran a simulation that is summarized in Figure 5. The results of the simulation are shown in Figure 14. Overall, the effects on pRF properties were much smaller than the changes we observed empirically although we simulated bias-induced response changes as much as 100% from leading to trailing edge. We found the bias to shift the pRF center in the direction of motion which is *incompatible* with our empirical results for low-contrast stimuli (but compatible with results for high-contrast stimuli). Furthermore, pRF size decreased for outward as compared to inward motion (compatible with empirical results), and gain, which is comparable to the signal change we reported, increased (incompatible with empirical results).

For all these reasons, we do not believe that the aperture-inward bias had a large effect on the pRF properties reported here - especially not on the eccentricity values for the low-contrast condition. This does not exclude the possibility, however, that it might have affected other pRF properties such as the pRF size estimation.

**Figure 5:**
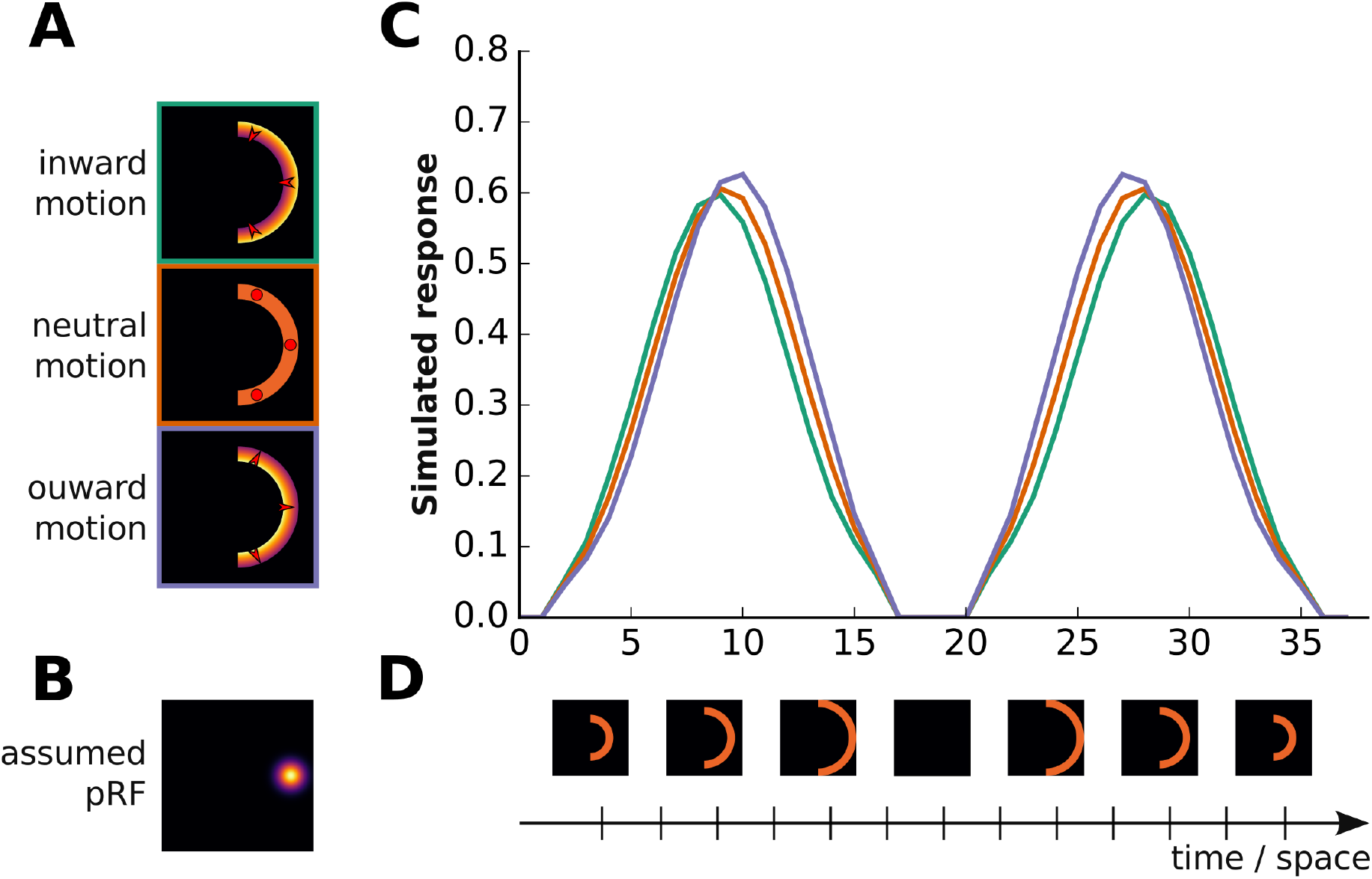
Simulation of the effects of aperture-inward bias on pRF response. When assuming higher responses to the trailing edge of motion, the pRF center is shifted in the direction of motion. **(A)** We simulated an aperture-inward bias by assuming higher responses to the trailing edge of motion. This implied higher responses to the outer aperture edge for inward motion (green), equal responses across the entire aperture for flicker motion (orange), and higher responses to the inner aperture edge for outward motion (violet). Red arrows indicate motion direction, red dots indicate absence of a particular motion direction. **(B)** We assumed a pRF that was centered on the right horizontal meridian with an eccentricity of 5.95°of visual angle and a pRF size of 1°(shown), 2°or 3°(not shown). **(D)** We assumed that a half-ring aperture first stepped from low (3.4°) to high (8.5°) eccentricity (15 steps, in 30s) and then, after some rest period (8s), stepped back from high to low eccentricities. **(C)** Responses were obtained by spatially convolving the assumed pRF with the different apertures presented in the simulation. Inward (green), flicker (orange) and outward (violet) motion yielded different responses, as shown by the three correspondingly coloured curves.

For the high-contrast data we observed pRF center shifts in the direction of motion, at least for mid-level visual areas. If we assume that these effects were mostly unaffected by aperture-inward bias, then how can we explain these shifts? One possible explanation is offered by the motion-direction asymmetry (MDA) model [12]. According to this model, neurons selective to inward motion should always show a spatial response profile that is higher in the inward part than the outward part of the receptive field, independently of which motion direction is presented. Conversely, outward-selective neurons should show higher responses in the outward part of the receptive field. A voxel that is assumed to show an averaged response profile of both inward- and outward-preferring neurons will then show an inward bias in case of inward motion and an outward bias for outward motion. In line with this model, we observed pRF shifts in the direction of motion for V3a/b and TO - areas that are known to contain a high proportion of motion-direction selective cells [66, 67].

If this explanation is correct, why did we not observe similar effects for the low-contrast data? Since we measured data for the same voxels, the same directional biases should have led to similar pRF shifts in the direction of motion for the low-contrast stimuli. Possibly, the effects did occur but were over-shadowed by the pRF shifts in the direction opposite to motion that underlie the perceptual shifts. This would explain why in the low-contrast condition for TO, where the effect of the hypothesized MDA effect was strongest, we no longer observed a shift in the direction opposite to motion.

Reversely, in LO where we did not find such an an MDA effect, the shift in the direction opposite to motion was very large.

We observed pRF shifts for the low-contrast stimulus in all visual field maps with the excpection of TO. This may be surprising as TO, which comprises human V5/hMT+, is most prominently known for its responses to motion and one might thus expect TO to show motion-dependent effects. There are (at least) two possible explanations for why we did not find pRF shifts in TO. First, it is possible that the MDA effect hypothesized by Liu [12] also affected the low-contrast data. For high-contrast stimuli, we observed the highest displacement in the direction of motion for area TO. It is thus possible that this effect counteracted the displacement of the pRF against the direction of motion in our measurements, and as a result, we found no eccentricity difference between inward and outward moving stimuli. Second, to link displacements in pRF to position shifts in perception, we assumed above that lower-level visual areas signal displaced position codes that are read-out by hierarchically higher populations. Given that TO is hierarchically the highest visual area included in our study, it is possible that TO represents a higher-order read-out area.

For both high-contrast and low-contrast pRF mapping data we found that inward motion evoked higher signal change than outward motion in early visual areas. This indicates that, at least up to the eccentricities probed in this study (maximum 8.5°), there might be a bias for motion towards the fovea (centripetal motion bias). Several fMRI studies [68, 69, 70] previously attempted to investigate radial motion biases but findings were confounded by aperture-inward bias. Wang et al. [13] found evidence for a weak centripetal motion bias in V1–V3 although they noted that their measured responses were dominated by the aperture-inward bias. Our results are fully consistent with the findings from Wang et al. [13] and offer additional empirical evidence for a centripetal motion bias in areas V1 to V3 without the complications of dominant aperture-inward bias.

Some fMRI studies also reported larger responses for motion towards the fovea in area hMT+ [71, 13], yet other studies did not find a difference in response amplitude between motion towards and away from the fovea [68, 72]. We observed a centripedal bias in TO for high-contrast stimuli but found no difference between inward and outward motion for low-contrast stimuli. Furture studies are therefore needed to further clarify radial biases in TO.

We consistently observed larger pRF size for inward compared to outward motion for both high- and low-contrast stimuli. We believe that several factors contributed to these differences. First, our simulations on the link between perceptual displacements and pRF shifts showed that the observed pRF size difference might contribute to the perceptual shift in the direction of motion. Second, our simulations on the influence of aperture-inward bias on pRF properties revealed that if aperture-inward bias were to affect our findings they would have resulted in pRF size difference in the direction observed here. Finally, Zeidman et al. [73] noted that for the pRF estimation procedure described in their paper, there was a high positive correlation between the model parameter for pRF size and the parameter for scaling the response. Given that we observed higher responses for inward as compared to outward motion, at least in early visual areas, this might have affected the estimation of pRF sizes. The expected effect is consistent with our results, with higher pRF sizes for inward motion than outward motion.

Although, we found no perceptual displacements at high stimulus contrast, we cannot conclude with certainty that position shifts are absent during high-contrast motion. It is possible that perceptual displacements were simply too small to be picked up by our current study design. For example, Fu et al. [10] found small displacements in the order of 0.03°to 0.10°for stimuli presented at 100% contrast. This suggests that perceptual displacements probably vary continuously with stimulus contrast and the exact relationship between stimulus contrast and perceptual displacements should be addressed in future studies by studying a wide range of contrast levels.

The choice to use checkerboard flicker in the current study, as opposed to counter-phased gratings, was motivated by two considerations. First, by setting the angular width of the checkerboard cubes appropriately, we could avoid that black and white elements are perceived to be displaced in radial direction for flicker reversals. By contrast, counter-phased gratings can give the impression of rings spontaneously “jumping” in radial direction when contrast reverses. Second, checkerboard flicker corresponds to the carrier pattern that is traditionally chosen for retinotopy [74] and pRF mapping [17]. It thus offered a direct reference point to other retinotopy and pRF mapping studies.

The choice of checkerboard flicker came with the downside that spatial frequency in the angular direction could not be matched to the inward and outward conditions. This usually did not have a visible impact on our results since the flicker condition tended both perceptually and in terms of the pRF properties to assume a position in-between the inward and outward conditions. However, there are two observations that might be explained by this design choice. First, at low contrast we observed that flicker elicited much higher responses than either of the other two conditions, at least in early visual areas. Second, at least for low-contrast stimuli, we observed that the slope for psychometric functions in the flicker condition was often flatter than for the other two conditions.

## 5 Supplementary material

**Figure 6:**
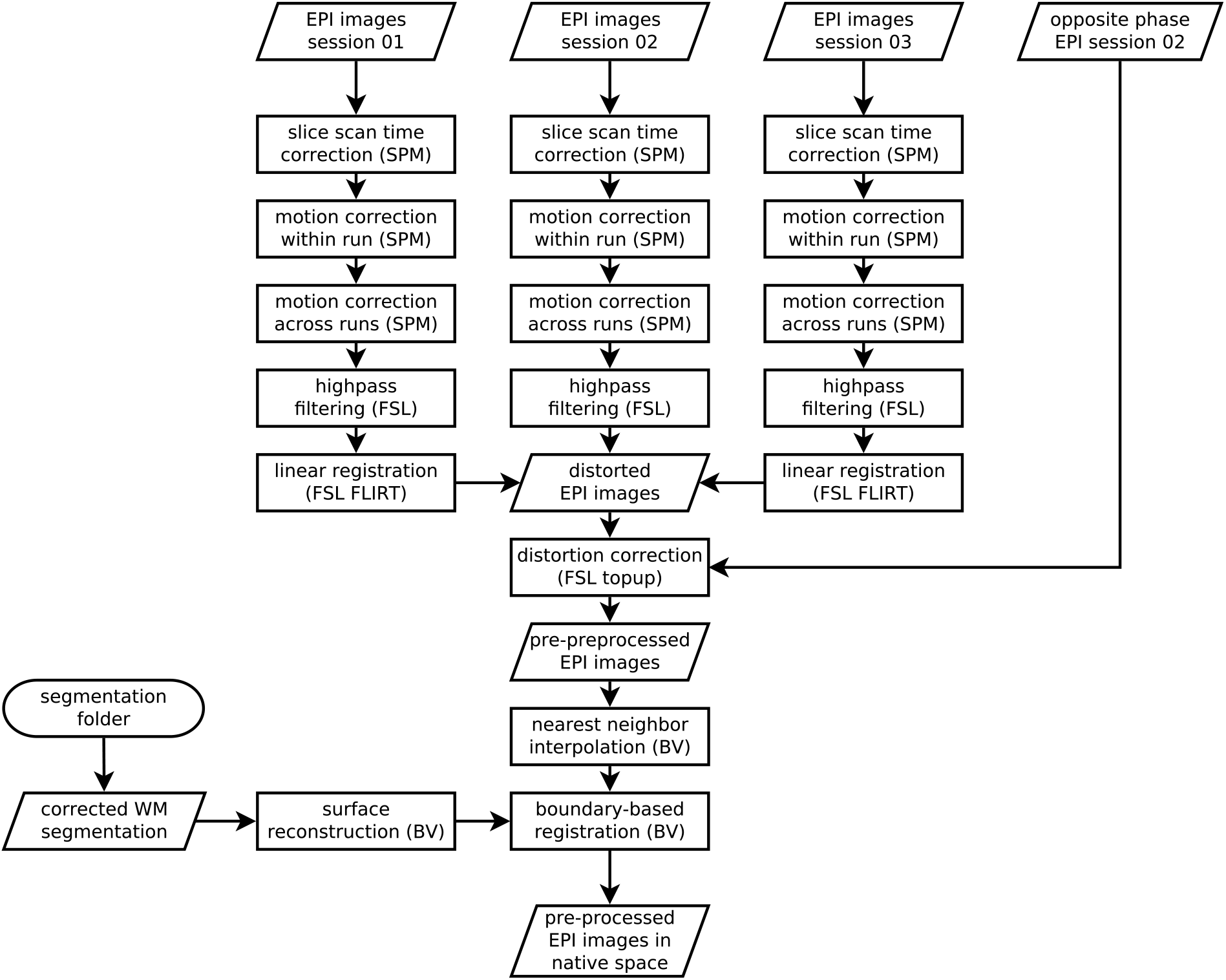
Overview of pre-processing pipeline for functional images. Rectangular shapes represent processing steps, rhombic shapes represent input or outputs and cylindrical shapes represent input or output locations.

**Figure 7:**
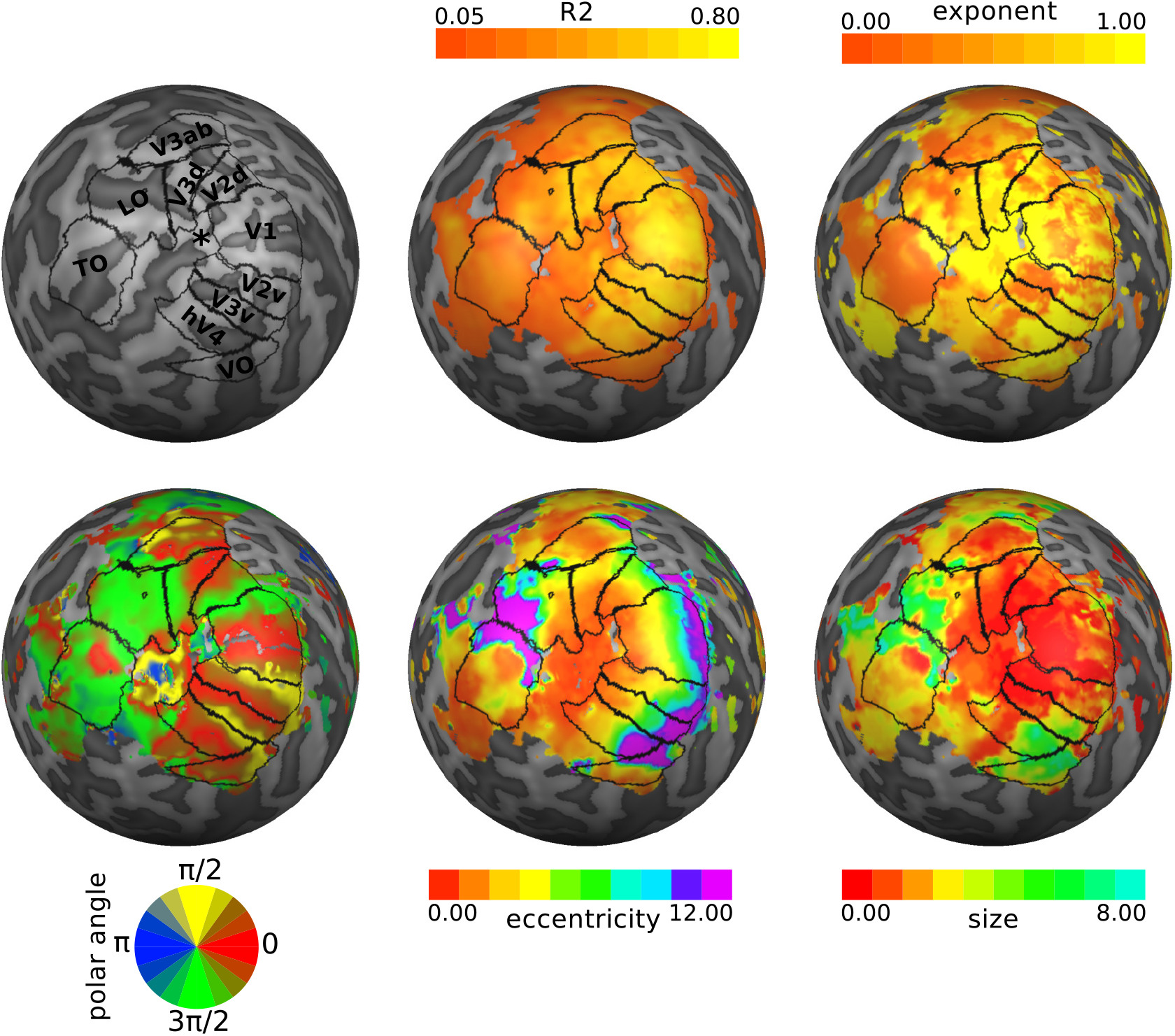
Overview of pRF mapping results used for delineation of visual field maps. The pRF properties are shown on the left hemisphere of an exemplary subject (S02). For better visualization the hemisphere reconstruction was morphed into a sphere. Dark-gray color indicates sulci and light-gray color indicates gyri.

**Figure 8:**
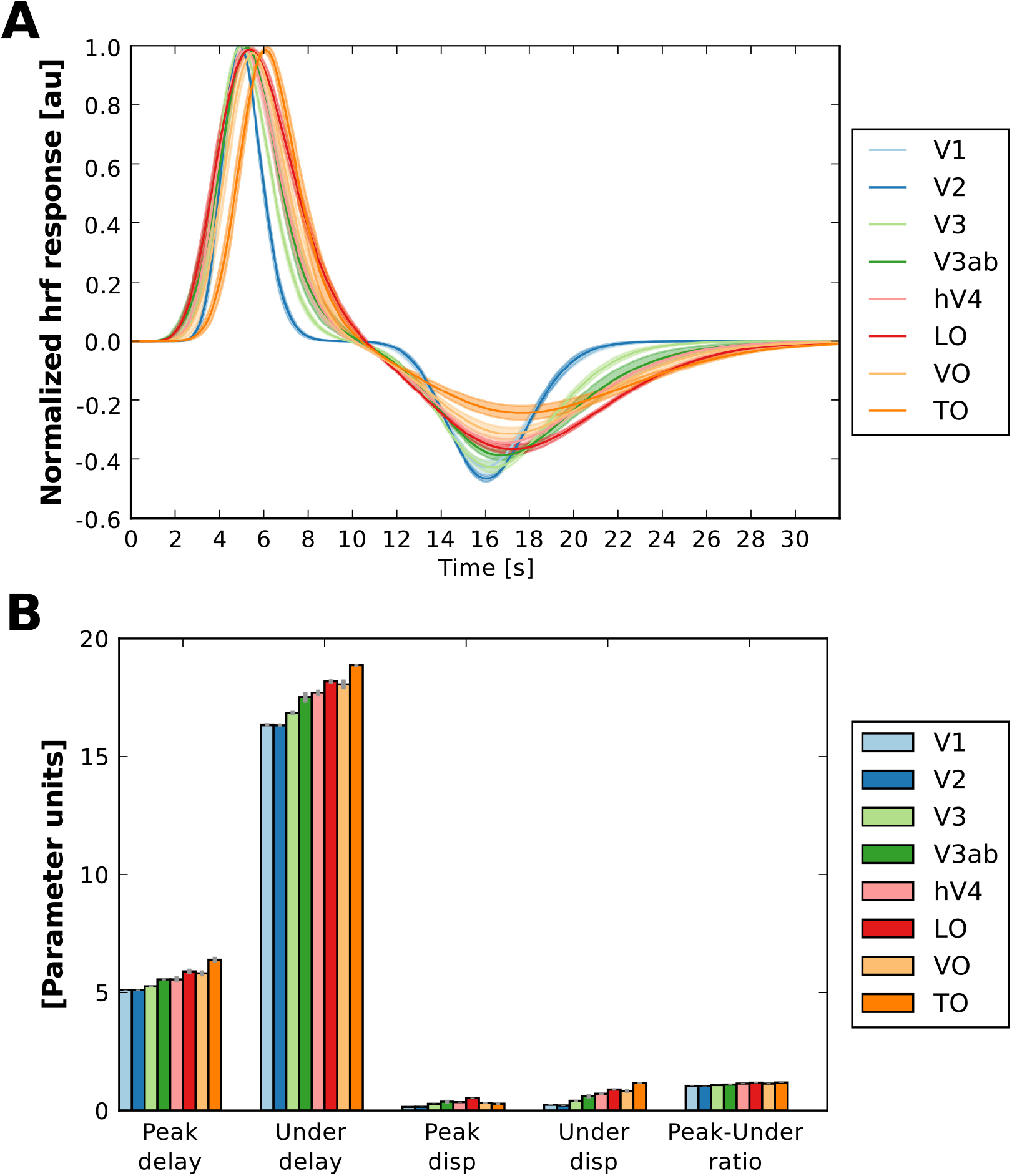
Fitted HRF parameters and hr functions. **(A)** Fitted hr functions. For visual display and improved comparison only, hr functions were normalized to set the peak for every HR function to 1. **(B)** Fitted HRF prameters. Shown are the averages across all subjects. Error shading and error bars indicate the standard error of the mean.

**Figure 9:**
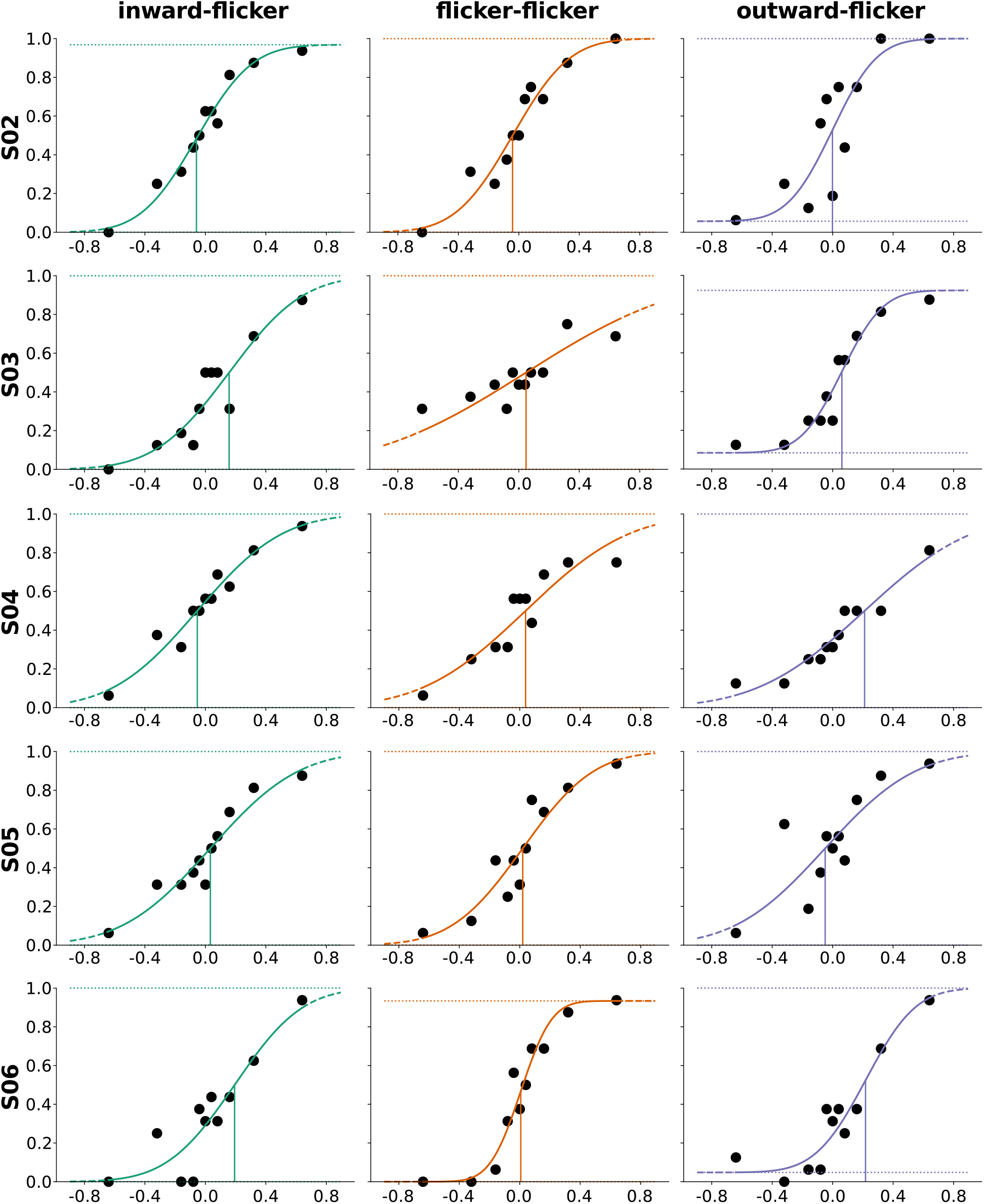
Single-subject psychometric data and functions for high contrast stimuli.

**Figure 10:**
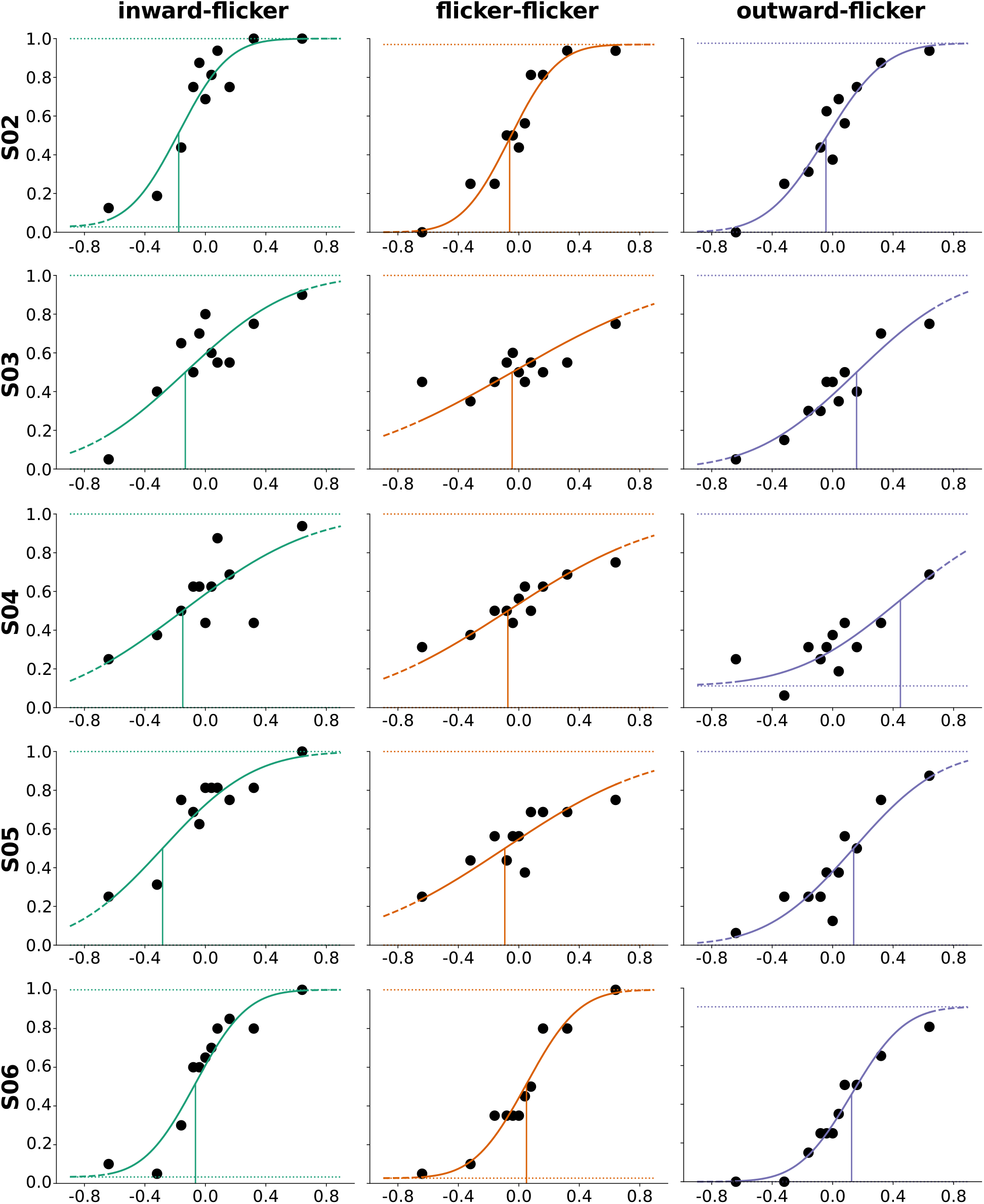
Single-subject psychometric data and functions for low contrast stimuli.

**Figure 11:**
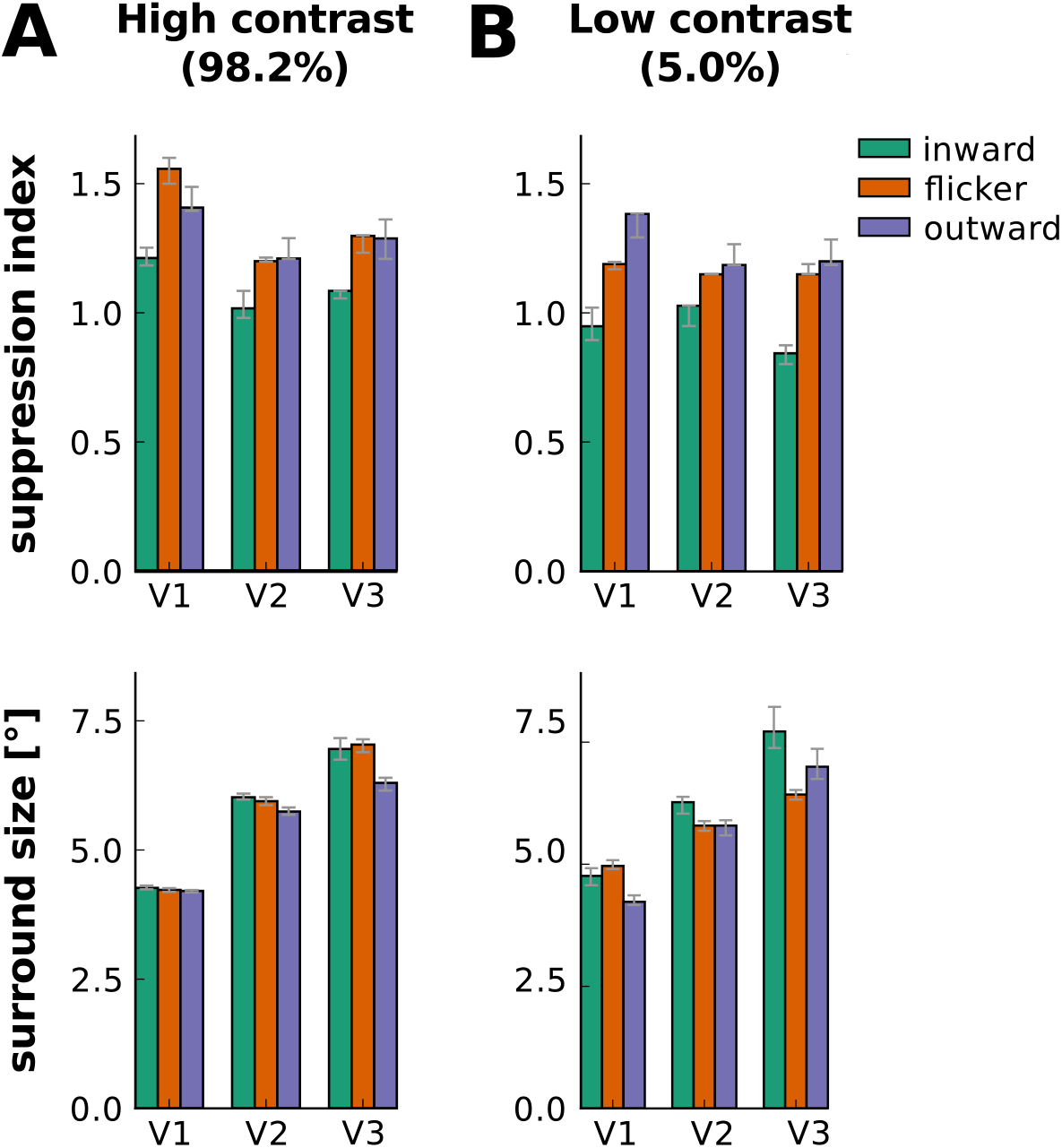
Effect of carrier motion on suppression index and suppressive surround size in early visual cortex. Same conventions as in Figure 3.

**Figure 12:**
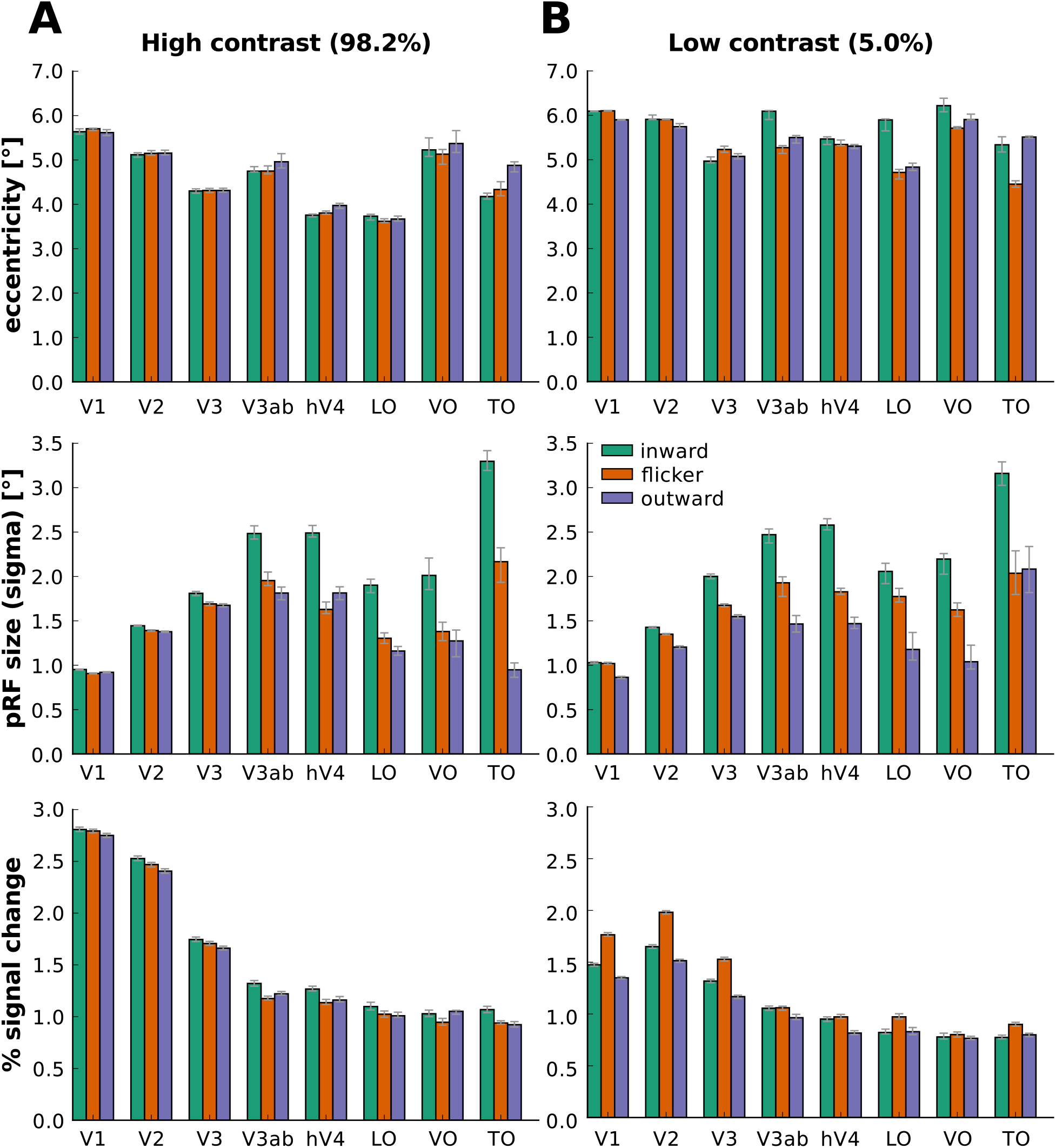
Effect of carrier motion on pRF properties when classical pRF model without suppressive surround is used. Same conventions as in Figure 3 with the difference that for pRF size the *σ* parameter of the 2D Gaussian is reported (in °of visual angle).

**Figure 13:**
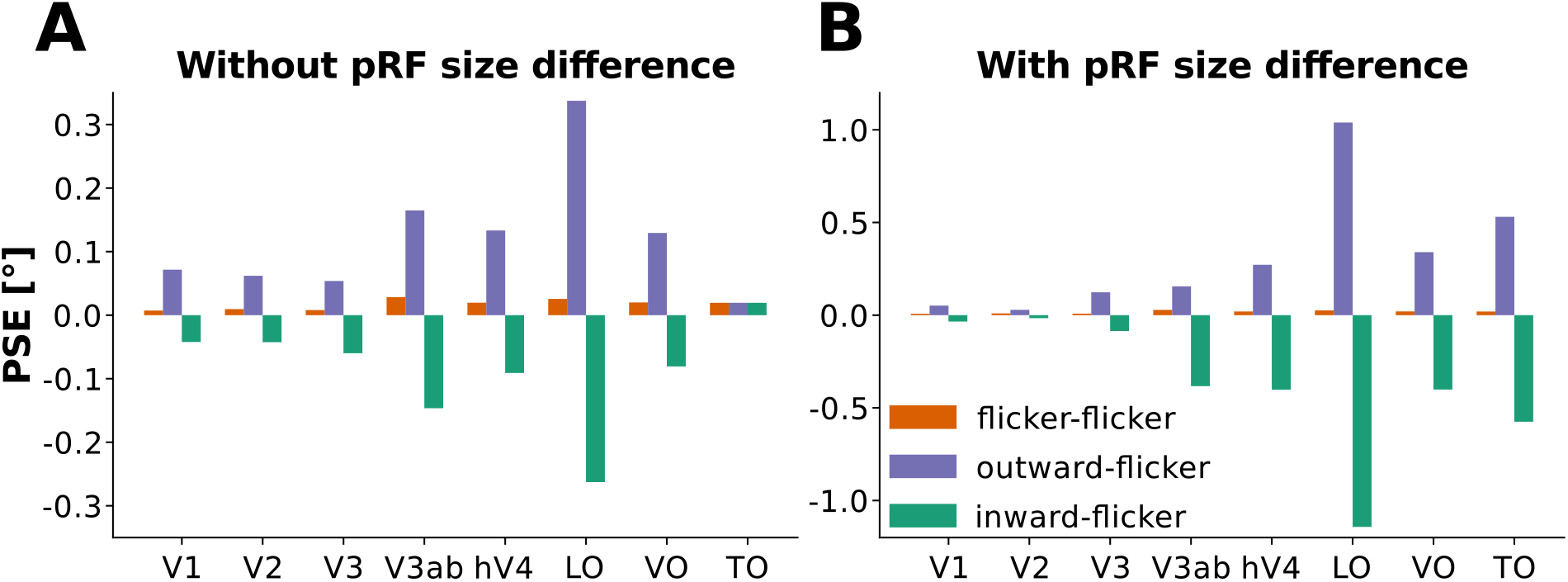
Simulated perceptual consequences of pRF shifts opposite to motion direction. A pRF center shift in the direction opposite to motion leads perceptual shifts in the direction of motion. **(A)** Point of subjective equality (PSE) for simulated psychophysics data. The three bar conditions represent simulated responses for the sequence of inward then flicker motion (green), flicker then flicker motion (orange) and outward then flicker motion (violet). We assumed that the eccentricity of the pRF shifted with the same magnitude that was observed empirically (see Figure 3, upper row). The pRF size was assumed to remain fixed. **(B)** PSE for simulated psychophysics data. Same conventions as in **(A)** but this time, in addition to shifting the pRF center, we also changed the pRF size in accordance with the pRF size difference that we observed empirically (see Figure 3, middle row). Note the different scale between the y-axes in **(A)** and **(B)**.

**Figure 14:**
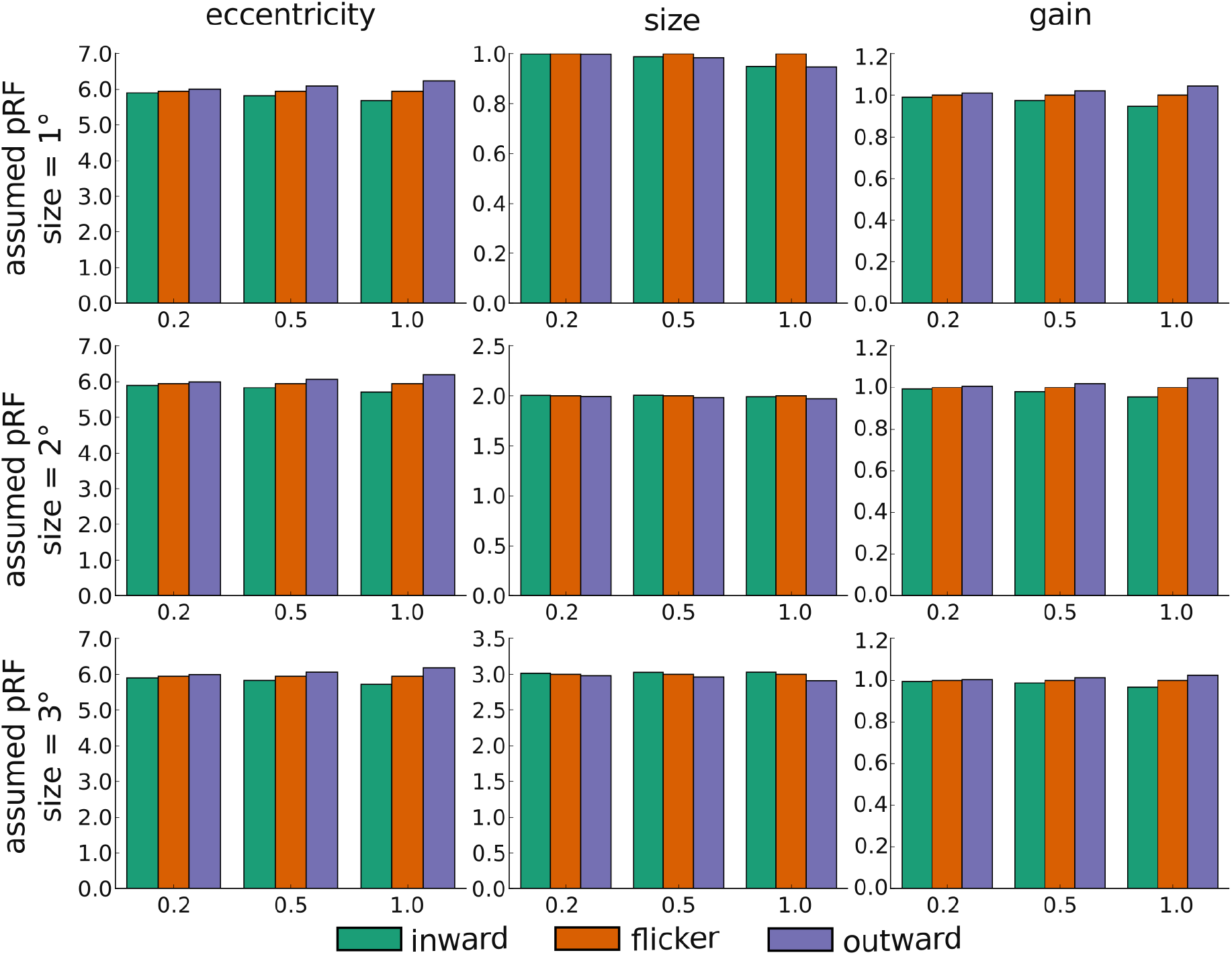
Simulated effects of aperture-inward bias on pRF response. We assumed a pRF that was centered on the right horizontal meridian with an eccentricity of 5.95°of visual angle. We varied the pRF size assuming either 1°, 2°or 3°, as shown in the different rows. The different columns show the effects of aperture-inward bias on different pRF properties: eccentricity, size and gain. The different colors represent the effect of inward motion (green), flicker motion (orange) or outward motion (violet). Different sections on the x-axis of each panel reflect the steepness of the modelled linear increase/decrease in response resulting from the aperture. The response was simulated to be 1 at the center of the aperture and we simulated an increase/decrease in response of either 0.2, 0.4, or 1.0 from aperture center to aperture edges.

## 6 Acknowledgments

We would like to thank Dimo Ivanov for help with adjusting the MP2RAGE sequence and Mario Senden and Brian Wandell for valuable advice.

